# An underlying bistability sets amplitude and explains temperature compensation in the cyanobacterial circadian clock

**DOI:** 10.64898/2026.05.03.722537

**Authors:** Yujia Liu, Gopal K. Pattanayak, Chris Chi, Helen Yoo, Aaron R. Dinner, Michael J. Rust

## Abstract

Circadian clocks create free-running biological rhythms with a period close to 24 hours. A universal property of these systems is temperature compensation, where the period of oscillation remains nearly invariant even as the amplitude changes with temperature. In the cyanobacterial system, where the core oscillator can be reconstituted from purified KaiABC proteins, we identify a key temperature-dependent positive feedback process: antagonism between KaiA and KaiB creates a bistable switch in protein-protein interaction. The region of bistability is strongly temperature dependent and correlated with oscillator amplitude. Combining this bistable mechanism with the temperature scaling of phosphorylation and dephosphorylation rates in a mathematical model recapitulates the overall temperature compensation of the oscillator. This capacity for history-dependent switching suggests that bistable dynamics underpin the generation of circadian rhythms.

**Significance Statement:** Circadian clocks are biological oscillators that anticipate the day-night cycle. The 24-hour time scale of the clock is remarkably insensitive to temperature owing to compensatory changes in amplitude and speed with temperature. In the cyanobacterial clock, we identify a bistability of the protein interaction network which determines oscillator amplitude and depends strongly on temperature, balancing the temperature dependence of dephosphorylation rates. This bistability indicates that oscillations arise from a combination of positive and negative feedback loops, a design principle used to maintain temperature compensation.

## Main text

A fundamental property of circadian rhythms is that the period of oscillation is nearly invariant to temperature changes in the physiological range (*1*). This phenomenon has been observed in all kingdoms of life and has posed a long-standing mechanistic puzzle: many biochemical processes, including enzymatic reactions and growth rates, are temperature dependent, typically accelerating by a factor of 2-3 with a 10 degree increase in temperature (*2*).

Rather than the system being completely temperature invariant, the amplitude of the oscillation—the range of chemical concentrations that occur during a cycle—compensates for changes in reaction rates (*3*). This results in an orbit in the space of biochemical concentrations that changes in size and location in response to temperature while the period remains nearly constant. The temperature dependence of the orbit shape is physiologically important, as it allows a circadian oscillator to respond to temperature changes in the environment by shifting phase (*4, 5*). Thus, a molecular understanding of temperature compensation must explain two features: the temperature-dependent scaling of the amplitude and the coincident scaling of reaction rates.

Remarkably, both of these features are present in the reconstituted phosphorylation oscillator created by the KaiABC proteins from cyanobacteria. Temperature compensation in this system was apparent even from its initial discovery (*6*), and more recent work clearly showed the temperature dependent scaling of amplitude of the phosphorylation rhythm (*7*). Because this system consists of only purified components, it presents the opportunity to identify the underlying mechanism of temperature compensation. However, achieving this goal requires isolating the processes that determine oscillator amplitude from those that determine period. Here we develop an experimental strategy to do so using partial reaction schemes that block one or more steps in the full oscillator.

In doing so, we identify a previously underappreciated positive feedback loop based on antagonism between the action of KaiA and KaiB that can create bistability in the KaiA-KaiB-KaiC interaction network in the absence of dynamic phosphorylation. We find that the region of bistability, defined by a range of KaiA concentrations that permit multiple steady states, is strongly temperature dependent. Combined with measurements of temperature-dependent phosphorylation and dephosphorylation rates, we integrate these observations to construct a mathematical model of the KaiABC oscillator that can recapitulate temperature compensation and temperature-dependent scaling of amplitude.

### The height of the KaiC phosphorylation peak scales with temperature

KaiC is a hexameric double-domain RecA-superfamily ATPase with the ability to autophosphorylate at two residues, Ser431 and Thr432, in the C-terminal, or CII, domain (*8*). Phosphorylation is stimulated by the action of KaiA which binds to the KaiC C-terminal tails and stimulates the catalytic cycle, in part by promoting nucleotide exchange (*9, 10*). Phosphorylation and dephosphorylation occurs at these two residues in a kinetically ordered sequence (*11, 12*).

Oscillation in the KaiABC reaction can be divided into two phases: a daytime phase when KaiA is active and promotes KaiC phosphorylation, and a nighttime phase when KaiA is inactive and KaiC phosphorylation decreases through autodephosphorylation (*13*). In the current understanding of the system, KaiA inhibition is dictated by the phosphorylation state of KaiC: phosphorylation on Ser431 enables KaiC to form KaiB•KaiC complexes. This process involves allosteric signaling from the phosphorylated CII domain to the N-terminal CI domain and requires ATP hydrolysis in CI to allow KaiB to bind (*14-16*). These KaiB•KaiC complexes then trap KaiA in an inactive conformation (*17*). When these complexes dissociate after dephosphorylation has occurred, active KaiA is released and the cycle begins anew.

To investigate how the phosphorylation cycle changes with temperature, we analyzed KaiABC reactions run at different temperatures using gel electrophoresis that can resolve specific phosphorylated states of KaiC (Fig. 1A). Consistent with previous reports, the period of oscillation is temperature compensated and the overall amplitude of the phosphorylation rhythm decreases as temperature is lowered (fig. S1) (*7*). Visualized in a plane defined by the phosphorylation of Ser431 and Thr432, we see that orbit size increase with temperature (Fig. 1A). This is consistent with the framework described above: the speed of the system as it traverses this orbit, determined by the phosphorylation and dephosphorylation rate constants, must increase to balance the larger orbit size at higher temperature (Fig. 1B and C).

**Fig. 1.**
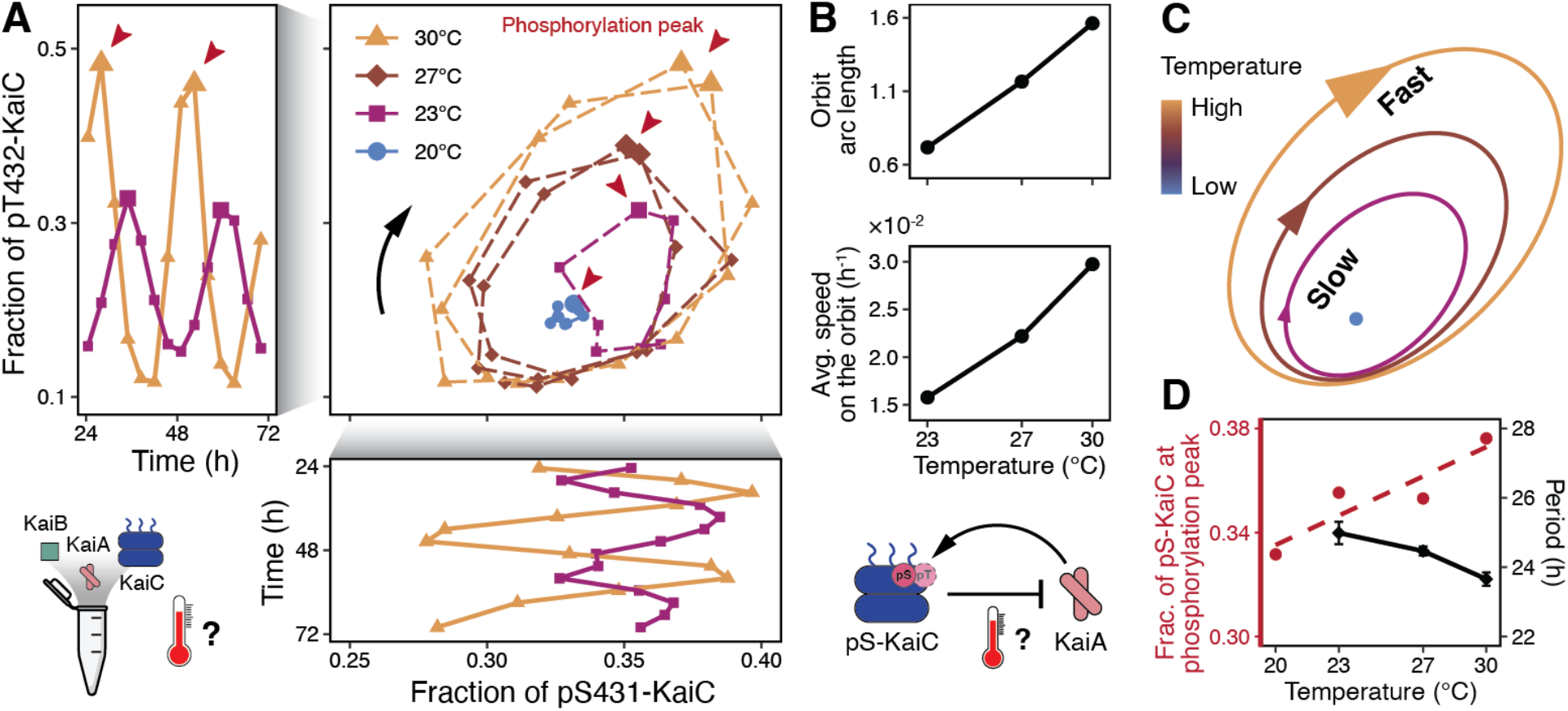
Oscillation orbit scales with temperature despite nearly invariant period. (**A**) Phosphorylation dynamics at two residues of KaiC, Ser431 and Thr432, in a reconstituted KaiABC reaction at different temperatures. The horizontal axis in the center panel represents the total fraction of KaiC phosphorylated at the Ser431 residue (pS431-KaiC), a sum of the fraction of KaiC phosphorylated only at Ser431 and both at Ser431 and Thr432. Similarly, the vertical axis represents the sum of the fraction of KaiC phosphorylated only at Thr432 and both at Ser431 and Thr432. Red arrows, peaks of total KaiC phosphorylation. Side panels, time series. Center panel, orbits in a phase space defined by KaiC phosphorylation. Stable oscillations at each temperature are plotted in the center panel: 23 °C, the 3^rd^ cycle recorded. 27 and 30 °C, the 2^nd^ and 3^rd^ cycles recorded. 20 °C, between 38.5 and 63 hours. (**B**) Temperature scaling of the orbit arc length and the average speed on the orbit as shown in (A). The orbit arc length is calculated by taking the sum of segment lengths the system traverses in the phase plane. The average speed is the arc length divided by the traversal time. (**C**) Schematic of temperature scaling of orbit size and reaction speed. The system travels faster on a larger orbit as temperature increases. (**D**) Summary of temperature dependence in the oscillator reaction. Red circles, fraction of Ser431 phosphorylated KaiC (the sum of those phosphorylated only at Ser431 and doubly phosphorylated at Ser431 and Thr432) at the total phosphorylation peak. Black diamonds, periods from fitting a sine function. Error bars, standard errors of the fitted periods.

A clue to the mechanism of how the orbit size changes with temperature comes from analyzing the pattern of phosphorylation in more detail. If inhibition of KaiA is simply proportional to the amount of Ser431-phosphorylated KaiC, we would expect the amount of phosphorylation needed to fully inhibit KaiA and initiate dephosphorylation to be independent of temperature. Instead, the data show that the composition of phosphorylation states at the turning points of the oscillation changes with temperature. In particular, the amount of the inhibitory Ser431 phosphorylation present at the phosphorylation peak—the transition between the phosphorylation (day) and dephosphorylation (night) phases—is higher at higher temperatures (Fig. 1D). This led us to investigate the possibility that the relationship between KaiC phosphorylation and KaiA inhibition is temperature dependent.

### An underlying bistability from KaiA-KaiB antagonism

To isolate the ability of KaiB•KaiC complexes to inhibit KaiA, we first adapted an assay to monitor KaiB•KaiC complex formation using a fluorescence polarization probe on KaiB (Fig. 2A). Previous work showed that formation of the KaiB•KaiC complex is slow and requires either phosphorylation on Ser431 or replacement of this residue with a phosphomimetic amino acid (*12, 15*). Here we used KaiC S431E;T432E (KaiC-EE) which binds KaiB efficiently and cannot change its phosphorylation state. When we combined KaiC-EE with KaiB at either 30 °C or 20 °C, we observed similar kinetics of complex assembly (0.12 vs. 0.10 µM^-1^ h^-1^) (fig. S2), suggesting that the intrinsic ability of phosphorylated KaiC to bind KaiB is not strongly temperature dependent.

**Fig. 2.**
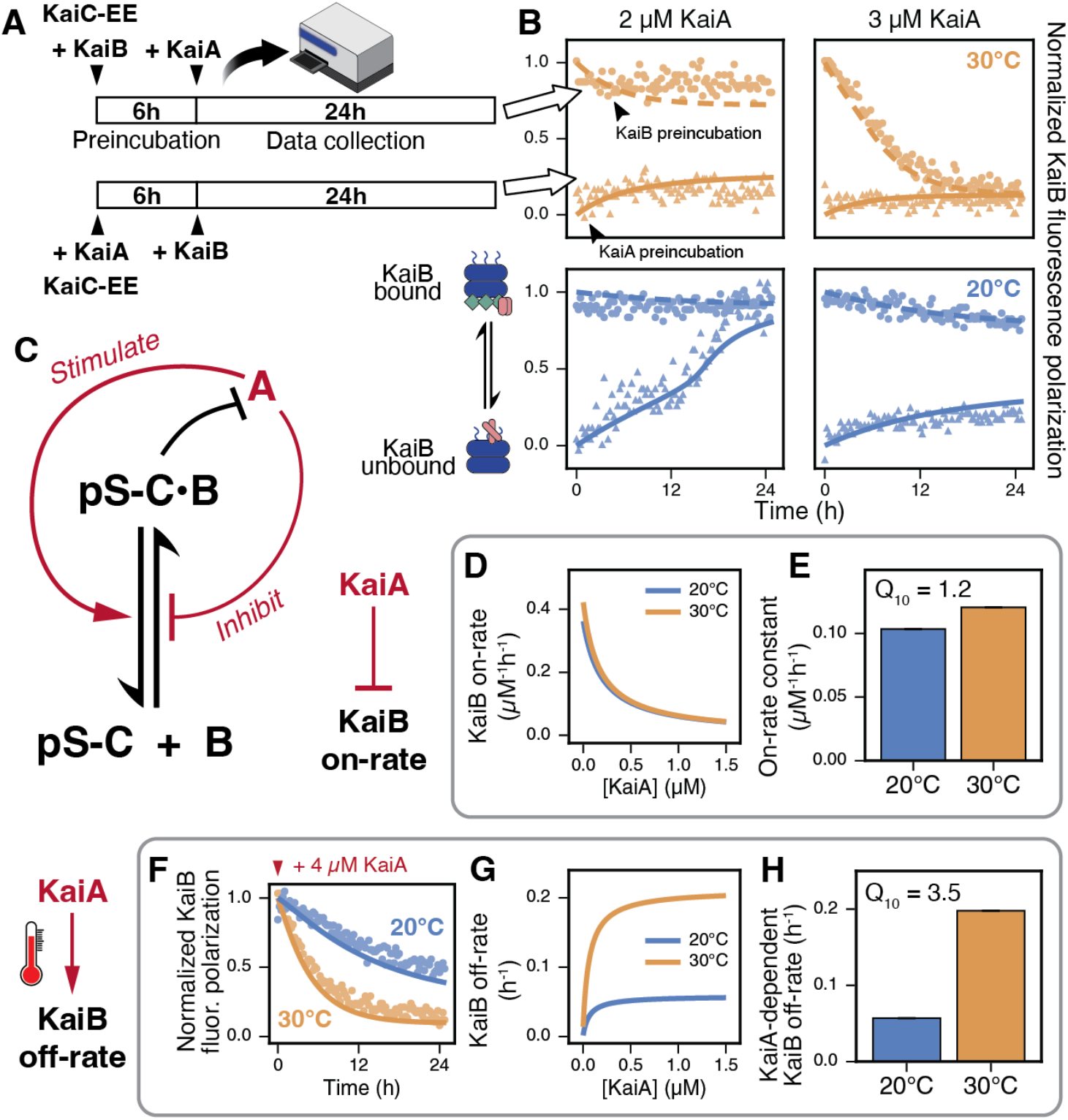
A temperature-dependent positive feedback loop in the KaiA–KaiB–KaiC interaction network. (**A**) Experimental protocol for probing KaiB•KaiC complex formation using fluorescence polarization. KaiC-EE was first preincubated with either KaiB or KaiA for 6 hours. Then, KaiA or KaiB was added to the respective reaction mixtures, such that the two conditions differ only in their order of addition. After protein addition, reaction mixtures were immediately moved to a microplate reader which records fluorescence polarization. (**B**) Normalized KaiB fluorescence polarization traces overlaid with model fit at selected KaiA concentrations. Circles and dashed lines, KaiC preincubated with KaiB. Triangles and solid lines, KaiC preincubated with KaiA. Columns are different KaiA concentrations used in the reaction and rows are different reaction temperatures. (**C**) Reaction scheme of the KaiA-KaiB-KaiC protein-protein interaction network. A, KaiA. B, KaiB. pS-C, KaiC in a Ser431-phosphorylated state. KaiA possesses the dual function of inhibiting KaiB•KaiC assembly and promoting KaiB release. (**D** and **E**) KaiB on-rate as a function of KaiA concentration and the fitted on-rate constant. Here we use Q_10_ to denote the ratio between the 30°C and 20°C value of a reaction rate constant. Error bars, standard errors of the fitted rate constants. (**F**) Dynamic change of pre-formed KaiB•KaiC complex probed by fluorescence polarization after adding 4 µM KaiA. Circles, experimental measurements. Solid lines, model fits. (**G** and **H**) KaiB off-rate as a function of KaiA concentration and the fitted off-rate constant. Error bars, standard errors of the fitted rate constants.

However, when we introduced KaiA into this assay, we observed two striking effects. First, at high KaiA concentrations, KaiB•KaiC complex assembly is blocked, even though this mutant KaiC has permanent phosphomimetic mutations that permit KaiB binding (fig. S2). This is consistent with our previous finding that KaiA acting on the KaiC C-terminus allosterically prevents KaiB binding (*18*). Indeed, we find that KaiA is unable to antagonize KaiB•KaiC complexes when the KaiC C-terminal tail has been deleted (fig. S3). Second, at intermediate KaiA concentrations, the outcome of the assay depends on the order of addition of the components. When KaiA is added first, KaiC appears to be locked into a KaiA-stimulated state that cannot bind KaiB. In contrast, when KaiB is added first, KaiB•KaiC complexes form that can then inhibit KaiA and prevent its action (Fig. 2B and fig. S2).

This history-dependence is the hallmark of systems that contains multiple stable steady states (*19*). We note that, even in the absence of phosphorylation dynamics, this system follows the motif of a double negative feedback loop: KaiB•KaiC complexes inhibit KaiA through stochiometric binding, but active KaiA signals allosterically through KaiC to inhibits KaiB•KaiC complex formation. Double negative feedback loops have an effective positive feedback logic and can give rise to bistability (*20*). Although the principles of thermodynamics prevent chemical reaction systems near equilibrium from exhibiting bistability, KaiB•KaiC complex formation is known to be a non-equilibrium binding process and requires ATP hydrolysis in the KaiC CI domain (*14*).

### The bistable region is strongly temperature dependent

When we repeated these experiments at 20 °C, we observed that the effect of KaiA on the system is clearly temperature dependent (Fig. 2B, fig. S2 and S4). At higher temperature, disassembly of existing KaiB•KaiC complexes occurs faster when KaiA is added, and the region over which bistable outcomes are possible shifts to lower KaiA concentrations. Overall, this suggests that, while KaiB•KaiC complex formation in the absence of KaiA is kinetically similar, KaiA’s ability to antagonize KaiB•KaiC is markedly stronger at higher temperatures.

To quantify the effect of active KaiA on KaiB•KaiC complexes we fit the kinetics of the order-of-addition preincubation experiments (Fig. 2B) to a bimolecular binding model where the free KaiA concentration non-cooperatively modulates assembly and disassembly rates (Fig. 2C and SI text). From the fit, we estimated the KaiB•KaiC on-rate and off-rate constants (Fig. 2E and H), which suggest that KaiA acts allosterically both to inhibit the complex assembly rate and to stimulate complex disassembly (Fig. 2D and G). This is reminiscent of the analysis of bistability in cell-cycle regulation where feedback to both forward and reverse processes broadly enhances the range of conditions supporting bistability (*20*). Notably, a previous study found that the bare KaiB•KaiC dissociation rate in the absence of KaiA is nearly temperature independent (*21*), while we find that the regulatory effect of KaiA on dissociation is highly temperature sensitive (Fig. 2F).

Bistable dynamics suggest an interpretation of the circadian oscillation as toggling between alternative metastable states that correspond to night and day. In the daytime state, KaiA is free and active. In the nighttime state, KaiA is inactive and bound in KaiA•KaiB•KaiC complexes with high fluorescence polarization. We used our bimolecular binding model to predict a bifurcation diagram showing regions of bistability visualized as steady-state abundance of KaiB•KaiC complexes vs. KaiA concentration (fig. S4). Importantly, the model predicts that the phosphorylation of KaiC should also be bistable as a function of KaiA concentration, separating the daytime and nighttime states of the system by a hysteretic switch (Fig. 3A).

**Fig. 3.**
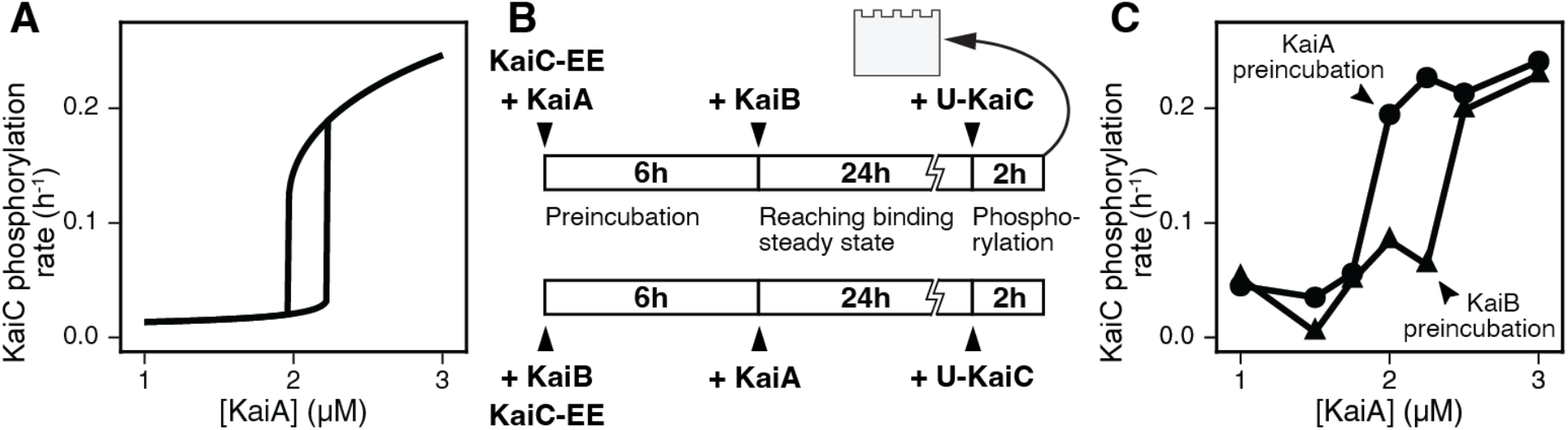
Positive feedback loop gives rise to a bistable switch. (**A**) Model-predicted KaiC phosphorylation rates at KaiB•KaiC binding steady-state as a function of KaiA concentration. The upper and lower branches of the bifurcation diagram can be elicited by initializing with low or high levels of KaiB•KaiC complex, respectively. Both KaiB and KaiC are set at 3.5 µM. (**B**) Experimental protocol for assaying the active KaiA amount in the reaction mixture using naïve unphosphorylated KaiC (U-KaiC) as a probe. Similar to the fluorescence polarization assay, KaiC-EE was preincubated with either KaiA or KaiB for 6 hours. KaiB or KaiA was subsequently added to respective reaction mixtures to create different orders of addition. Reactions were then incubated for 24 hours to reach binding steady state before probe U-KaiC was added. Two-hour end-point samples were collected and analyzed by SDS-PAGE. (**C**) Experimentally determined bifurcation diagram showing initial KaiC phosphorylation rates as a function of KaiA concentration. The upper and lower branches were elicited by preincubating KaiC-EE with KaiA or KaiB, respectively. Each data point is an average of duplicate measurements.

To test this prediction, we developed an assay to measure the residual KaiA activity in these reactions. After preincubation and allowing Kai protein complexes to form, we introduce unphosphorylated wildtype KaiC. This “probe” KaiC will then autophosphorylate at a rate that depends on the available, non-sequestered KaiA (Fig. 3B). We collected time courses to monitor KaiC phosphorylation rates, confirming that phosphorylation activity depends sensitively on the amount of KaiA in the reaction and exhibits a temperature-dependent bistable region (Fig. 3C and fig. S5).

### Dephosphorylation of Ser431 accelerates with temperature

Intuitively, we expect the amplitude of oscillation to depend on the extent of the bistable region (Figure 3A). That is, as the level of inhibitory (Ser431 phosphorylated) KaiC increases throughout the day, the system should initially remain close to the upper branch of the diagram where KaiA activity prevents KaiB•KaiC complex assembly. Only when the system approaches the bifurcation point, where the upper branch disappears, can KaiB•KaiC complexes form, allowing the dephosphorylation phase to begin, causing the system to traverse the lower branch.

As shown in Figure 1, reduction of amplitude requires a slower average reaction rate along the phosphorylation cycle to maintain a 24-hour period. Thus, we sought to characterize phosphorylation and dephosphorylation rates at different temperatures. We measured phosphorylation kinetics at each site at 30 °C vs. 20 °C and their dependence on KaiA concentration (fig. S6). Fitting a model of KaiC phosphorylation and dephosphorylation network (fig. S7) to the data, we find that the phosphorylation rate for the Thr432 site has a mild temperature sensitivity, increasing by a factor of 1.6 between 20 °C and 30 °C, and a mild increase by a factor of 1.4 in the concentration of KaiA needed for half-maximal activation (fig. S6 and table S1).

In contrast, when we allowed KaiC to dephosphorylate in the absence of KaiA (Fig. 4A-B, fig. S8), we found a striking temperature dependence, particularly in the dephosphorylation of the inhibitory site Ser431 (Fig. 4B), increasing by about 6-fold from 20°C to 30°C. Consistent with this estimate, dephosphorylation is also markedly temperature dependent in single site mutants (fig. S9). Although early reports showed that both KaiC phosphorylation and dephosphorylation kinetics were nearly temperature invariant, these studies had shorter time courses and did not resolve modifications on each site (*6*). Since Ser431 is only dephosphorylated after Thr432, these temperature dependent kinetics are likely only apparent on longer timescales (fig. S8).

**Fig. 4.**
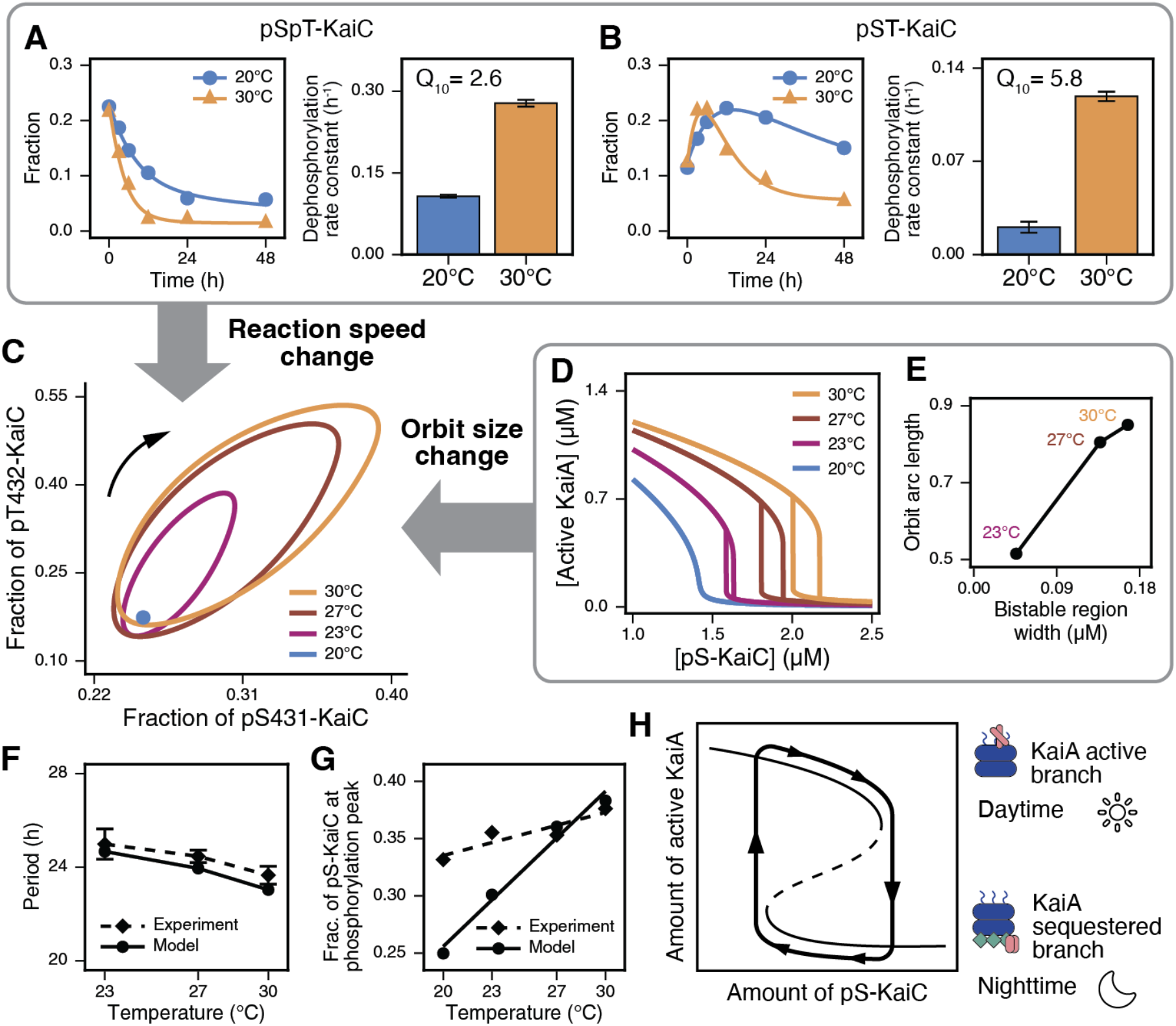
A balance between reaction speed and bistable region changes explains temperature compensation. (**A** and **B**) Site-specific KaiC dephosphorylation kinetics in the absence of KaiA, and their fitted rate constants. Here we use Q_10_ to denote the ratio between the 30°C and 20°C value of a reaction rate constant. Dots, experimental measurements. Solid lines, model fits. Error bars, standard errors of the fitted rate constants. (**C**) Predicted stable limit cycles or steady states plotted in a phase plane of the two KaiC phosphorylation sites using a model incorporating KaiA–KaiB antagonistic mechanisms. (**D**) Prediction of steady-state levels of active KaiA at different amounts of Ser431 phosphorylated KaiC (pS-KaiC) using the bimolecular binding model. The upper and lower branches can be elicited by initializing with either low or high levels of KaiB•KaiC complex. (**E**) Orbit arc lengths in (C) plotted against the bistable region widths in (D). (**F** and **G**) Comparison between model-predicted (circles and solid lines) and experimentally determined (diamonds and dashed lines) period (F) or fraction of Ser431 phosphorylated KaiC at the phosphorylation peak (G). Error bars, standard errors of the periods from fitting experimental data. (**H**) Schematic interpreting sustained oscillation as traversing the distinct branches set by un underlying bistability. In the daytime, the system traverses the upper branch high in active KaiA, which allows KaiC phosphorylation to drive the system rightwards. At the cusp determined by the bistability, the system is forced to jump to the lower branch, starting the nighttime. Light solid and dashed lines, stable and unstable steady states of the underlying bistability.

### A balance between changes in bistability and reaction speed explains temperature compensation

To understand how these temperature-dependent biochemical processes contribute to the oscillatory dynamics of the system, we constructed a mathematical model (fig. S10 and SI text). We modified an existing ordinary differential equation model that produces circadian oscillations based on rates measured at 30 °C (*14*). We first introduced temperature scaling of the phosphorylation and dephosphorylation rates using rate constants measured at 20 and 30 °C (Fig 4A-B and table S1). In stark contrast to experimental observations, the period markedly lengthens as temperature decreases whereas the amplitude remains stable (fig. S11).

We then sought to introduce the temperature dependent antagonism between KaiA and KaiB. Some existing mathematical models of the KaiABC oscillator lack such a mechanism (*14*), and those with sufficient expressivity do not operate in a parameter regime capable of reproducing the bistability we observe (*18, 22*). Since a major temperature-dependent phenomenon we observed in our measurements is a KaiA-stimulated complex disassembly rate, we reformulated the structure of the Phong et al. model (*14*) to include both KaiA-dependent on-rates and disassembly rates (fig. S10). Model parameters are constrained by simultaneously fitting the biomolecular binding subnetwork model to the KaiB•KaiC complex formation data (Fig. 2), and the full model to the KaiC phosphorylation oscillation data (Fig. 1) at different temperatures (table S2, see Supplementary Text for further details).

The temperature scaling of rates in this full model predicts a similar bistable effect to that seen in experiments, now visualized by the presence of alternative stable active KaiA states at a given phosphorylation level (Fig. 4D). Combined with the temperature dependence of the phosphorylation and dephosphorylation rates, this model reproduces temperature compensated circadian rhythms, but only when both temperature dependent effects are present (Fig. 4F and fig. S12). Plotting orbits in the phase space of the two phosphorylation sites (Fig. 4C), we see that the orbits mirror the scaling and shifting of the bistable regions (Fig. 4D-E), consistent with the interpretation that circadian oscillation tracks the underlying bistability. The qualitative trends seen in the data are also correctly recapitulated by the model: the amplitude of the phosphorylation rhythm increases with temperature (fig. S12), and the amount of phospho-Ser431 KaiC at the peak scales with temperature (Fig. 4G).

## Discussion

Many features of the Kai oscillator are remarkably temperature independent. In this study, we sought to identify temperature-dependent processes that are needed to explain the decrease in oscillator amplitude and ultimate failure when temperature is decreased. In this analysis, we identified a previously under-studied double-negative feedback loop where KaiA can act allosterically on KaiC to oppose its own inhibition. This creates a positive feedback interaction that is strong enough to create bistability, where either a KaiA-active or KaiA-inactive state can be stably maintained when phosphorylation is fixed.

A balance between a temperature-dependent KaiA feedback loop and temperature-dependent KaiC dephosphorylation is sufficient to achieve a compensated period, as shown in our mathematical model. Previous studies argue against a delicate balance model with fine-tuned temperature dependencies across the entire biochemical network, in part based on an observation that many period-altering mutants also preserve temperature compensation (*5, 23*). Here, by measuring isolated biochemical processes, we discovered that temperature effects primarily impact a sparse set of steps in the oscillator cycle. These temperature-dependent steps are, importantly, involved in both positive feedback loops (KaiA opposes its own inhibition) and negative feedback loops (KaiC dephosphorylation relieves KaiA inhibition). These effects are in balance to allow amplitude to change while period remains nearly constant.

General arguments about temperature compensation mechanisms have concluded that there must be some rate constants whose increase causes period elongation to balance an inverse dependence on other rate constants (*24*). Here we can identify the rate constant describing KaiA-stimulated dissociation of KaiB•KaiC complexes as falling into this class and directly observe the mechanism: increasing the width of an underlying bistable region, which would lengthen the period in the absence of balancing acceleration of phosphorylation and dephosphorylation rates (fig. S12).

What is ultimately the structural origin of the temperature-dependent bistability? To activate KaiC phosphorylation, KaiA binds to the KaiC C-terminus (*25*), inducing conformational changes proximal to the phosphorylation sites in CII. In contrast, the ability of KaiA to disrupt KaiB•KaiC complexes must involve distal signaling from the KaiA binding site on CII to the KaiB binding site on CI. We show that the ability of KaiA to act proximally to drive phosphorylation in CII is quite temperature invariant (fig. S6), while the distal effect on CI is temperature sensitive. Since the ATPase rates of the KaiC domains are themselves insensitive (*26, 27*), this points to the allosteric signaling between the CII and CI domains itself being a point of temperature sensitivity. Consistent with this picture, the known mutations in KaiC that break temperature compensation, such as Arg393, lie near the interface of the two domains (*28, 29*).

Though circadian oscillators are often understood as delayed negative feedback loops, our finding that bistable dynamics appear when the phosphorylation state is frozen, indicates that a negative feedback-only picture cannot be qualitatively correct: bistability implies the existence of a strong positive feedback loop. The importance of positive feedback loops is clear in other biological oscillators such as the eukaryotic cell cycle, where positive feedback at the post-translational level has the potential to create bistability that then becomes a free-running oscillator when coupled to protein synthesis and degradation (*30*). The close relationship between bistability and oscillation can also be seen in simple models such as the FitzHugh-Nagumo equations modeling neural excitability (*31*). These dynamical principles may be conserved in circadian clocks generally: a phosphoswitch between antagonizing motifs has been identified in a critical slowdown in the degradation kinetics of Period2, a core mammalian clock protein, indicative of an underlying positive feedback loop (*32*). Notably, the critical slowdown disappears at a lower temperature, reminiscent of the temperature dependence we observe in KaiB•KaiC complex formation kinetics in the presence of KaiA. This raises the intriguing possibility that temperature-modulated positive feedback interactions may underlie temperature compensated circadian oscillators across kingdoms, despite the divergence of their molecular components.

The KaiA-dependent positive feedback loop we elucidate here is likely at the heart of sustained oscillations in the KaiABC reaction. Though phosphorylation is required for oscillation, we reinterpret the mechanism of oscillation as one where the changing level of Ser431 phosphorylation moves the system along an underlying bifurcation diagram arising from protein-protein interaction. This picture can intuitively explain the otherwise mysterious recent discovery of sustained temperature-compensated oscillations in a single-site mutant of KaiC (T432V) possessing only the Ser431 site (*16*). We propose that the daytime and nighttime phases of the system are fundamentally defined by the hysteretic branches of an underlying bistability. These self-reinforcing dynamical states are then slowly tuned by protein phosphorylation, resulting in oscillation (Fig. 4H). The existence of an underlying bistability may point towards the evolutionary path that led to self-sustaining rhythms. A bistable system based on positive feedback could allow a cell to retain a memory of a daytime or nighttime state that would persist in the face of transient environmental fluctuations. The introduction of negative feedback through the regulation of phosphorylation would allow this switch to become an oscillator, as in the modern cyanobacterial system.

## Acknowledgments

We thank Rust Lab members for useful discussions. We thank Mingxu Fang, Tian Yang, and Rob Rodriguez for their help with protein purification. We thank Carrie Partch for the gift of sortase.

## Author contributions

Conceptualization: CC, ARD, GKP, YL, MJR

Methodology: CC, ARD, GKP, YL, MJR, HY

Investigation: CC, ARD, GKP, YL, MJR, HY

Supervision: ARD, MJR

Writing: CC, ARD, GKP, YL, MJR

## Data, code, and materials availability

Code used to generate mathematical predictions is available on Zenodo. All data are available in the main text or the supplementary materials.

## Supplementary Materials

Materials and Methods

Supplementary Text

Figs. S1 to S12

Tables S1 to S2

## Materials and Methods

### Recombinant protein expression and purification

#### KaiA

KaiA was expressed as an N-terminal glutathione S-transferase (GST) fusion protein using vector plasmid pGEX6P-1 in *Escherichia coli* strain DH5α. Expression cultures were set up by diluting a dense overnight starter culture into two 1 L LB media with 50µg/mL carbenicillin. Expression cultures were incubated with shaking at 37°C until the optical density (OD) at 600 nm reaches 0.6, when the cultures were induced by 100 µM of isopropyl β-D-1-thiogalactopyranoside (IPTG) and allowed to express the recombinant protein at 16°C overnight with shaking. Cells were harvested the next morning by centrifugation and flash-frozen in liquid nitrogen.

To purify KaiA, we first resuspended the cell pellets in a lysis buffer with 1mM adenosine triphosphate (ATP) and lysed them using an Emulsiflex-C3 homogenizer (Avestin). We then passed the clarified supernatant through a 1 mL GSTrap HP column (Cytiva) using an Äkta go liquid chromatography system. We washed the column and cleaved the tag on-column by injecting 160 U PreScission Protease (Cytiva) and incubating at 4°C overnight. The eluate was further purified using a ResourceQ anion exchange column (Cytiva) before we buffer-exchanged it into Kai protein storage buffer (20 mM Tris-HCl pH 8.0, 150 mM NaCl, 5 mM MgCl2, 0.5 mM ethylenediaminetetraacetic acid (EDTA), 1 mM ATP, and 10% v/v glycerol) using a 10 kDa MWCO Amicon centrifugal filter (Millipore). The resulting KaiA solution was flash-frozen and stored in small aliquots.

#### KaiB

KaiB was expressed and purified as described previously (*14*).

#### KaiC

KaiC was expressed as an N-terminal small ubiquitin-like modifier (SUMO) fusion protein using a pET-28b(+) plasmid vector, with a Strep-6xHis dual tag N-terminally fused to the SUMO moiety and a FLAG tag fused downstream of SUMO, in *E. coli* strain BL21(DE3). Expression culture was prepared by diluting a dense overnight starter culture into 1 L LB media with 12.5 µg/mL kanamycin. The expression culture was incubated with shaking at 37°C until the OD at 600 nm reaches 0.4, when the culture was induced with 200 µM IPTG and allowed to express at 20°C overnight with shaking. Cells were harvested the next morning by centrifugation and immediately resuspended with lysis buffer (50 mM Tris-HCl pH 8.0, 150 mM NaCl, 1 mM MgCl_2_, 5% v/v glycerol, and 1 mM ATP). We then lysed the cells by passing through an Emulsiflex-C3 homogenizer. We added 100 units of Benzonase (Sigma-Aldrich) during the lysis process to reduce viscosity created by nucleic acids within the cell. After clarifying the lysate using centrifugation, we flowed the supernatant through 1 mL Strep-Tactin XT resin (IBA Lifesciences) packed in a 15 mL gravity-flow column. The column was then sequentially washed with a low-salt buffer (50 mM Tris-HCl pH 8.0, 150 mM NaCl, 1 mM MgCl_2_, 5% v/v glycerol, and 1 mM ATP), a high-salt buffer (low-salt buffer supplemented with 350 mM NaCl), and again low-salt buffer to re-equilibrate. KaiC was eluted using a buffer containing 50 mM biotin (IBA Lifesciences). To cleave the expression tags (Strep-6xHis-SUMO. FLAG tag is preserved), 4 µM homemade SUMO protease Ulp1 was added to the eluate, which was then incubated at 4°C overnight. Cleaved KaiC was further purified using a Sephacryl S-300 HiPrep size-exclusion column (Cytiva) and simultaneously buffer-exchanged into the aforementioned Kai storage buffer. The resulting KaiC solution was flash-frozen and stored in small aliquots.

#### KaiC S431E;T432E

KaiC S431E;T432E (KaiC-EE) was expressed and purified as described previously (*14*).

### In vitro KaiABC oscillation assay

A 64 µL oscillator reaction was assembled containing 1.5 µM KaiA, 3.5 µM KaiB, and 3.5 µM KaiC in reaction buffer (20 mM Tris-HCl pH 8.0, 150 mM NaCl, 5 mM MgCl_2_, 0.5 mM EDTA, and 1 mM ATP) in a twin.tec LoBind 96-well plate (Eppendorf), which minimizes absorption of biomolecules. The reaction mixture was incubated at the target temperature using a GEN2 thermocycler module on the deck of an OT-2 liquid handling robot (Opentrons). The robot was programmed to take 2 µL samples and quench in 10 µL sample buffer (62.5 mM Tris-HCl pH 6.8, 2% w/v sodium dodecyl sulfate (SDS), 0.05% w/v bromophenol blue, 10% v/v glycerol, and 5% v/v β-mercaptoethanol) every 3.5 hours up to 73.5 hours. The sampling interval of 3.5 hours was chosen not to be a divisor of the 24-hour period, thereby providing denser phase coverage over multiple days of measurement.

The samples were analyzed on a 10% sodium dodecyl sulfate-polyacrylamide electrophoresis (SDS-PAGE) gel (375 mM Tris-HCl pH 8.8, 10% v/v acrylamide/bis, 0.1% w/v SDS, 0.05% w/v ammonium persulfate (APS), and 0.5‰ v/v tetramethyl ethylenediamine (TEMED)), which resolves different KaiC phosphorylation states, and subjected to SYPRO Ruby staining (Invitrogen). Analysis of the fluorescent band intensity was performed using ImageJ software.

### KaiB•KaiC formation fluorescence polarization assay

#### KaiB fluorescence labeling

KaiB was fluorescently labeled following a previously described sortase-mediated N-terminal ligation protocol (*33*). Recombinantly expressed KaiB with an N-terminal glycyl residue, as a result of the cleavage of an N-terminal HRV-3C site, was exchanged into a sortase buffer (50 mM Tris-HCl pH 7.5, 150 mM NaCl, and 10 mM CaCl_2_) using 7 kDa MWCO Zeba spin desalting columns (Thermo Scientific). A 2 mL reaction was assembled containing approximately 50 µM KaiB, 5µM 6xHis-sortase, and a fluorescein-labeled peptide, FAM-LPETGG, then incubated at 4°C overnight protected from light. We passed the reaction mixture through a 1 mL HisTrap HP column (Cytiva) the next day using an Äkta go system to remove sortase. Relevant flow-through fractions were further purified using a Sephacryl S-300 HiPrep size-exclusion column and finally concentrated using 10 kDa MWCO Amicon centrifugal filters until the protein solution became visibly yellow. The resulting labeled KaiB solution was flash-frozen and stored in small aliquots. Sortase and FAM-LPETGG were gifts from Carrie Partch.

#### Fluorescence polarization assay

We set up the reaction by mixing 3.5 µM KaiC-EE with (a) 3.5 µM KaiB and 1% v/v FAM-KaiB stock solution or (b) different concentrations of KaiA in the reaction buffer and incubated it at the target temperature for 6 hours (preincubation). Then, either (a) the target concentration of KaiA or (b) 3.5 µM KaiB and 1% v/v FAM-KaiB stock solution were added before the reactions were moved to a 384-well non-binding black microplate (Corning) and monitored by a Spark microplate reader (Tecan) using the fluorescence polarization program (excitation wavelength = 485 nm, emission wavelength = 535 nm, monochromator) for 25 hours.

### KaiA activity assay

The reaction was set up as described previously for the fluorescence polarization assay where KaiB and KaiA were added in different orders. Unphosphorylated KaiC (U-KaiC) was freshly prepared by incubating the freezer-stock KaiC with 5 mM ATP at 30°C for 30 hours. After a 6-hour preincubation and a 24-hour second-step incubation, 3.5 µM of U-KaiC was added to the reaction mixture and 2-hour end point samples were taken. Samples from each condition containing approximately 100 ng of KaiC probe were subjected to SDS-PAGE analysis using a homemade 10% polyacrylamide gel. Electrophoresis was performed for 5.5 hours for improved resolution between different phosphorylation states of wildtype KaiC and KaiC-EE. The gel was then stained with SYPRO Ruby and analyzed using ImageJ. The initial phosphorylation rate of probe KaiC over 2 hours is estimated by dividing the change in the phosphorylation fraction between the end point and the start point by the time duration.

### Model fitting

#### KaiC phosphorylation and dephosphorylation processes

KaiC phosphorylation data were acquired by incubating 3.5 µM U-KaiC with different concentrations of KaiA at the target temperature and taking samples during a 6-hour time course. KaiC dephosphorylation data were acquired by incubating 3.5 µM freezer-stock KaiC at the target temperature and taking samples during a 48-hour time course. The samples were analyzed on a homemade 10% polyacrylamide gel, stained with SYPRO Ruby, and quantified using ImageJ.

To quantify the phosphorylation or dephosphorylation rates of individual steps, we fit the KaiC phosphorylation and dephosphorylation network model as described in **Fig. S7** to the data by minimizing a loss function defined as follows:

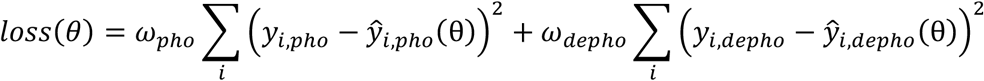

Where *θ* denotes a set of parameters. *y*_*i*_ denotes an experimental measurement and 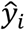 denotes a corresponding model prediction. *pho* represents the phosphorylation kinetics data and *depho* represents the dephosphorylation kinetics data. We manually set the weights as ω_*pho*_ = 0.5 and ω_*depho*_ = 1 to account for a larger number of phosphorylation datapoints.

We performed the minimization using simulated annealing as a first step (SAMIN solver) and the interior-point Newton method as a second step (IPNewton solver) using the Optim.jl package (Julia). Standard error of the i^th^ parameter, θ_*i*_, was calculated using

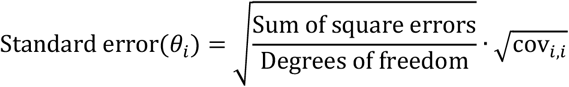

Where the degrees of freedom is the number of data points minus the number of parameters in the model. The covariance matrix was estimated by calculating the inverse of the Hessian matrix of the loss function evaluated at the best-fit set of parameters (ForwardDiff.jl).

#### KaiB–KaiC bimolecular binding model

The bimolecular binding model, described in detail in the Supplementary Text, was fit to fluorescence polarization measurements of KaiB•KaiC complex formation. Prior to model fitting, the fluorescence polarization data were normalized to account for dependence on the instrument and experimental conditions, including KaiA concentration. Baselines at each KaiA concentration were established by averaging a group (N = 4, manually picked) of the lowest and highest polarization values. The polarization data were normalized by linearly scaling the low and high baselines to 0 and 1, respectively. Time-series of the concentration of KaiB•KaiC complex were predicted by the model, initiated from either no complex or a steady-state binding level in the absence of KaiA. Similarly, model predictions were normalized by linearly scaling the two initial concentrations of KaiB•KaiC complex to 0 and 1, where the high initial concentration represents the upper bound of complex formation because of the antagonizing effect of KaiA.

Parameters of the bimolecular binding model, including those describing modulation of the assembly and disassembly rates by KaiA, were inferred by simultaneously fitting the order-of-addition fluorescence polarization data and the KaiC phosphorylation oscillation data. The oscillation data were included to constrain the absolute level of KaiB•KaiC complex formation predicted by the bimolecular binding model, which cannot be inferred from fluorescence polarization measurements alone, and to accommodate the full oscillator model, which is based on an existing model that lacks the underlying bistability. The parameters were estimated by minimizing misfit between experimental data and model predictions, expressed as a loss function:

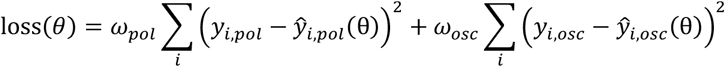

Where θ represents a set of parameters, *y*_*i*_ represents an experimental measurement, and 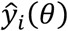 represents a corresponding model prediction given the parameters, and ω represents a weight applied to each dataset: *pol* denotes the fluorescence polarization measurements, and *osc* denotes the KaiC phosphorylation oscillation measurements. Here ω_*pol*_ = 1 and ω_*osc*_ = 30. Weights were chosen by hand to compensate for the much higher density of data points in the fluorescence polarization data sets. Parameters in the oscillator model describing phosphorylation and dephosphorylation were fixed during model fitting. We performed the optimization and calculated the standard errors of the parameter using the same procedure as for fitting the KaiC phosphorylation and dephosphorylation data.

## Supplementary Text

### KaiB–KaiC bimolecular binding model

A positive feedback loop model recapitulates dynamics of KaiB•KaiC complex formation in the presence of KaiA, where KaiA stimulates KaiB off-rate and inhibits KaiB on-rate, together antagonizing its own sequestration by the KaiB•KaiC inhibitory complex (**Fig. 2C**). Formally, we describe the action of KaiA not sequestered by the KaiB•KaiC complex (free KaiA, denoted as *A*_*free*_) on the on- and off-rates of KaiB by Michaelis-Menten kinetics:

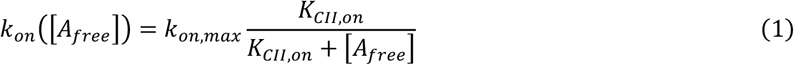

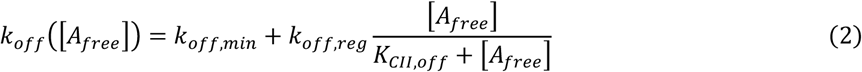

The dynamics of the system is then simply given by the law of mass action:

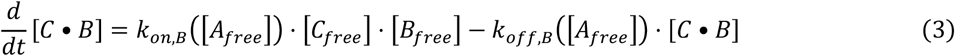

Where *C* denotes a KaiB-binding compatible KaiC species. *B* represents KaiB. In experimental settings, we used the KaiC-EE mutant which mimics a KaiC state permanently phosphorylated at both Ser431 and Thr432 residues.

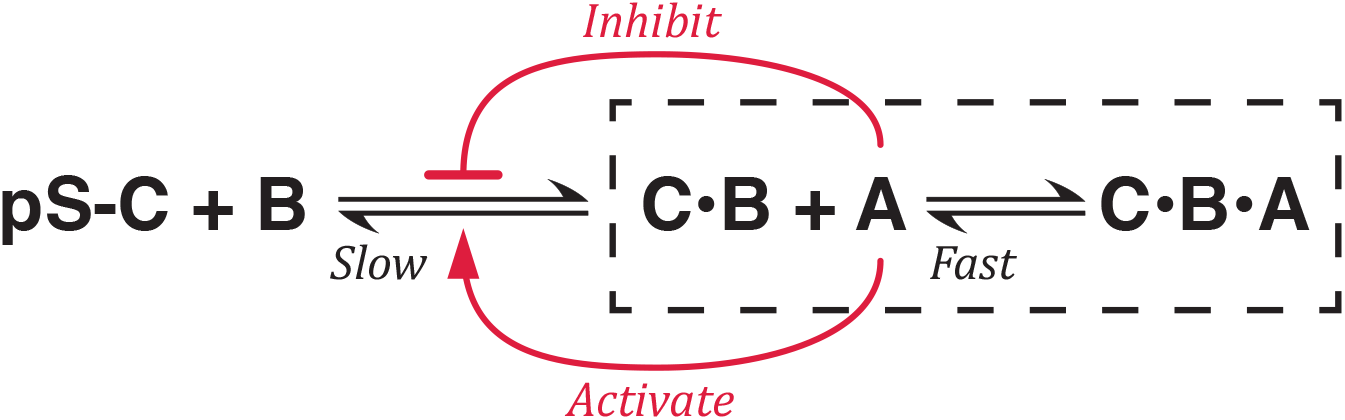

With conservation law, we have

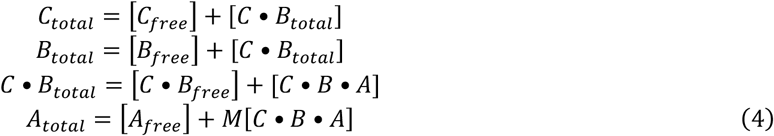

Where *M* is a stoichiometric constant. To determine the amount of *A*_*free*_, we take advantage of the fact that KaiA sequestration kinetics is much faster than that of KaiB•KaiC complex formation (diagram of reaction scheme above) and assume that the capture and release of KaiA by KaiB•KaiC is always at a steady state. Then, such processes are governed by an equilibrium constant

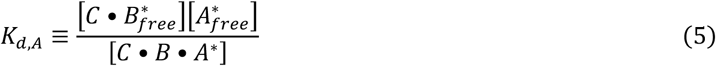

Where the asterisk signifies a steady state. Solving **Eq. 5** and **4** together gives us

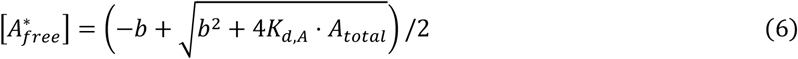

Where

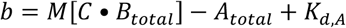

Note that since now free KaiA concentration is expressed as a function of KaiB•KaiC concentration, KaiB•KaiC formation can be fully described by a one-dimensional ODE, **Eq. 3**.

### Oscillator model

We constructed the full oscillator model (**fig. S10**) by incorporating KaiA-stimulated dissociation of KaiB•KaiC complex and KaiA-inhibited KaiB–KaiC assembly into an existing model described by Phong et al. (model A) (*14*). The new model is a six-dimensional ODE:

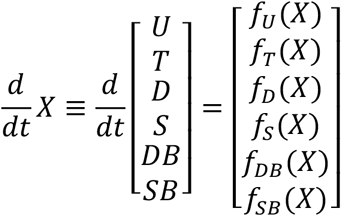

Where *U, T, D, S, DB*, and *SB* are shorthand for different phosphorylation states and binding states of KaiC. *U*, <u>u</u>nphosphorylated KaiC. *T*, KaiC phosphorylated at <u>T</u>hr432 residue only. *D*, KaiC <u>d</u>oubly phosphorylated at Ser431 and Thr432. *S*, KaiC phosphorylated at <u>S</u>er431 only. *DB*, D-KaiC in complex with KaiB. *SB*, S-KaiC in complex with KaiB. Now, the terms governing KaiB•KaiC assembly and disassembly are exactly given by **Eq. 3**, the bimolecular binding model. The full model describes both the assembly and disassembly of the KaiB•KaiC complex and the phosphorylation and dephosphorylation of KaiC. For simplicity, only the equations governing the *DB* and *SB* states are given below, involving KaiB•KaiC formation. All other equations are as described in (*14*).

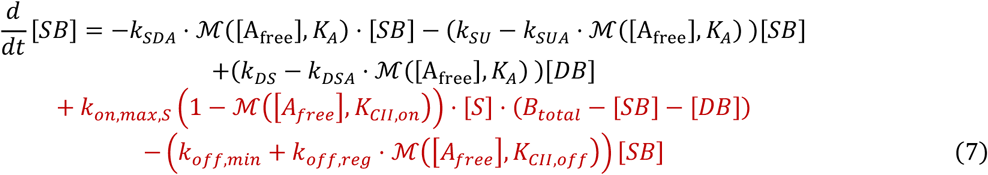

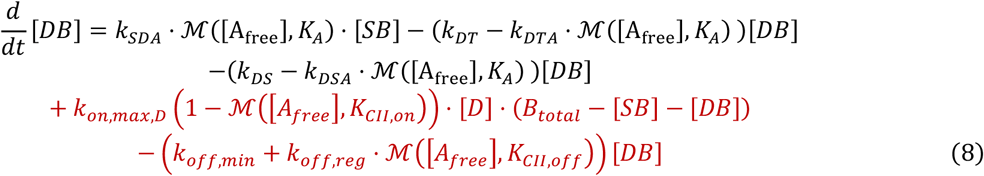

Where the free KaiA concentration is given by

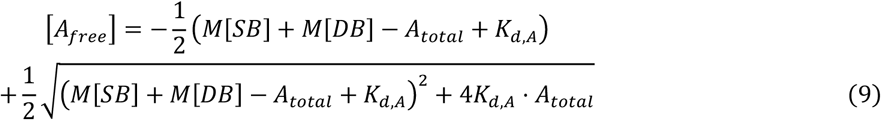

And ℳ is a Michaelis-Menten function,

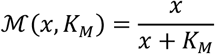

The naming convention for phosphorylation or dephosphorylation rate constants is as follows: the subscript denotes first the reactant and then the product, in some followed by an *A* to signify KaiA-dependence of the process. E.g., *k*_*SDA*_ is the reaction rate constant describing the *S* → *D* phosphorylation process, which is KaiA-dependent.

The terms describing KaiB•KaiC formation are newly introduced into the existing model and highlighted in red. Note that conservation laws dictate that the equations governing *D* and *S* also deviate from the existing model due to the changes in *DB* and *SB* equations.

In this work, when numerically solving the ODE model, we use total concentrations [*KaiA*] = 1 μ*M* and [*KaiB*] = [*KaiC*] = 3.5 μ*M*. A set of parameters describing the KaiABC system at 30 °C is given by a combination of those described in (*14*) (phosphorylation and dephosphorylation) and **table S2** (KaiB•KaiC formation). The rate constant describing KaiB and S-KaiC assembly (*k*_*on*_,_*max,S*_) is assumed to be 1.5 times higher than that for D-KaiC, following the observations in (*18*) in principle. Our implementation of the model is published in Zenodo (*34*).

### Introduction of temperature effects into the oscillator model

Leveraging biochemical assays where certain steps in the KaiABC reaction network are blocked, we discovered that temperature affects two sets of biochemical processes: bistability and KaiC phosphorylation and dephosphorylation. Phosphorylation and dephosphorylation rate constants at 20°C and 30°C were independently estimated by fitting the model in **fig. S7** to experimental data measured at these temperatures. The fold change of a parameter at 30 °C with respect to its 20 °C value is then calculated, named Q_10_ (since it is a fold change over 10 degrees of temperature change) (**table S1**). Temperature effects on the phosphorylation and dephosphorylation rates were then introduced into the model by fixing the original set of parameters from (*14*), which describes the 30°C behavior, and calculating the parameters of the system at 20°C by scaling using the corresponding Q_10_ coefficients. Incorporating the fitted parameters indirectly minimizes the modifications introduced to the original model. During parameter estimation, two phosphorylation steps, *U → SpT* and *U → pST*, were well constrained and, therefore, only the Q_10_ associated with those steps were included in the full oscillator model. Temperature scaling of three dephosphorylation steps were selected to be incorporated: *SpT → U, pSpT → pST*, and *pST → U*. Other dephosphorylation steps and phosphorylation steps were modeled as temperature-independent in a parsimonious fashion.

Temperature scaling of bistability was modeled by directly adopting the parameters of the temperature-dependent KaiB–KaiC bimolecular binding model (**Fig. 2C** and **table S2**), which were fitted using experimental data acquired at 20°C and 30°C. The parameter describing the system at an in-between temperature, e.g., 27°C, was calculated by linear interpolation of the 20 °C and 30 °C parameter sets.

---

**Fig. S1.**
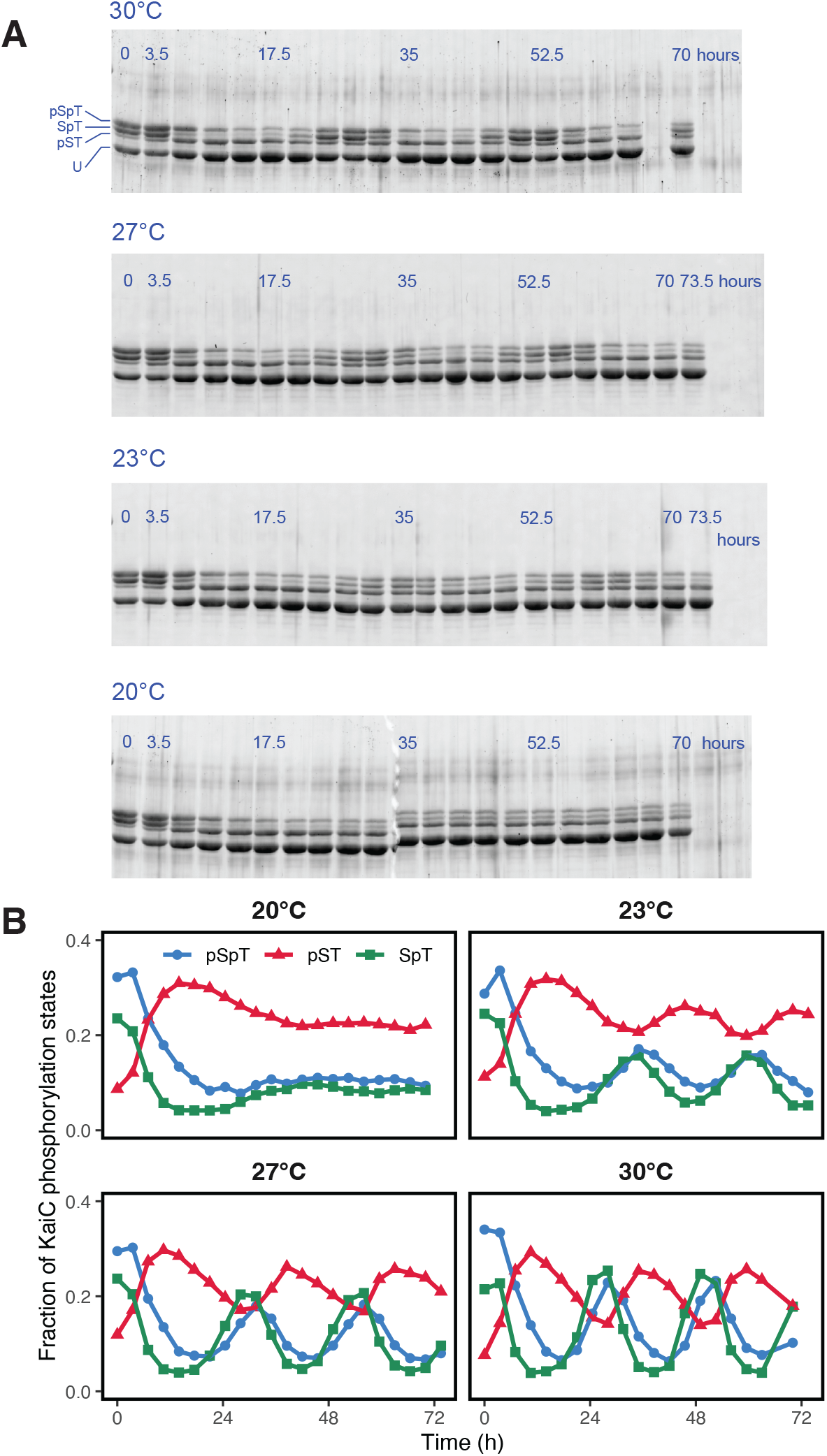
KaiABC reaction using purified components run at different temperatures. (**A**) Time course samples, collected every 3.5 hours, were analyzed by SDS-PAGE to resolve distinct KaiC phosphorylation states at two residues, Ser432 and Thr432. The four bands from top to bottom are doubly phosphorylated (pSpT), phosphorylated only at Thr432 (SpT), phosphorylated only at Ser432 (pST), and unphosphorylated (U). (**B**) Fractions of KaiC phosphorylation states, determined by quantifying gel band intensity.

**Fig. S2.**
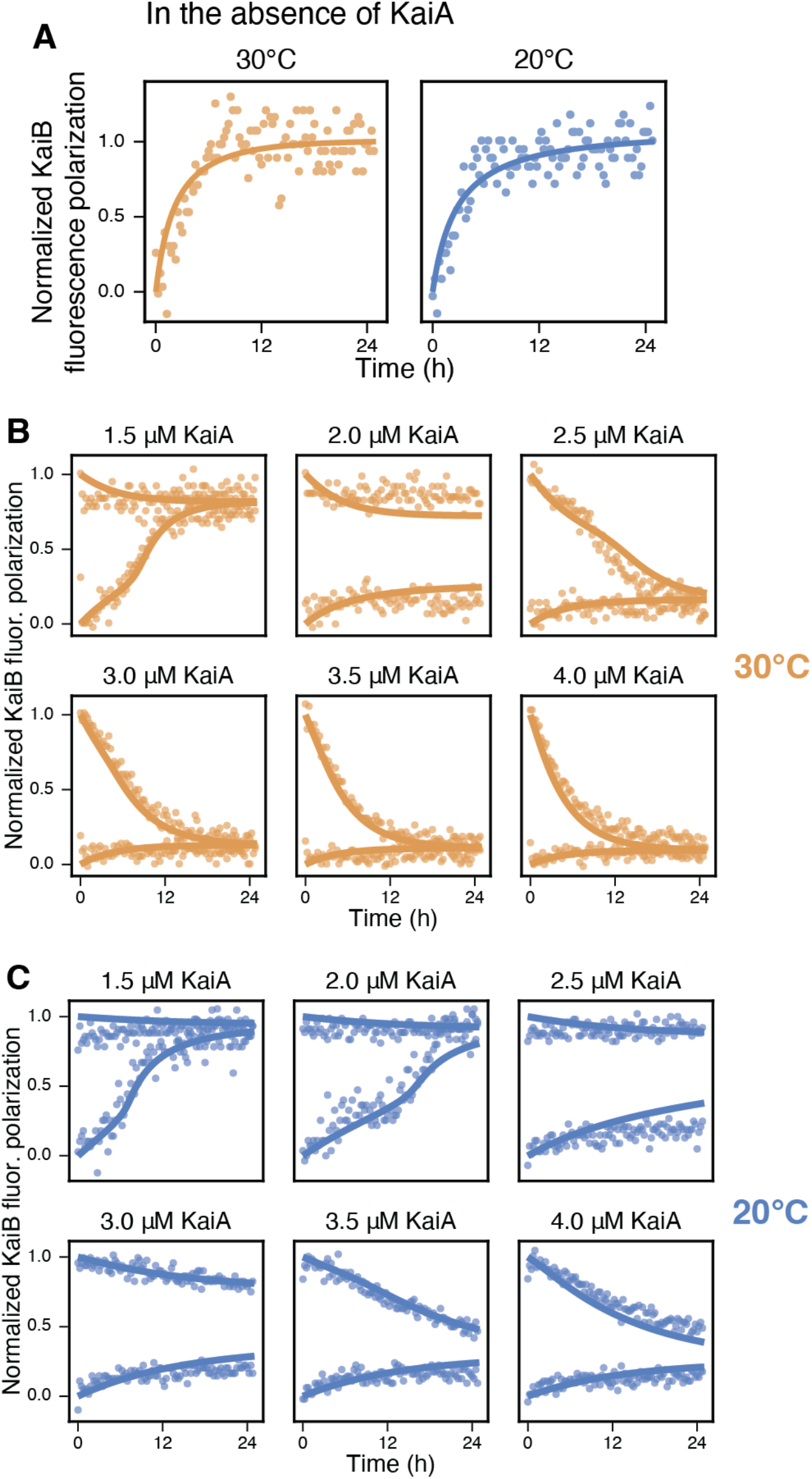
KaiB–KaiC-EE binding kinetics measured by polarization of a KaiB-attached fluorescence probe (**A**) in the absence of KaiA or (**B-C**) supplemented with different concentrations of KaiA. Circles, experimental measurements. Solid lines, fits using a bimolecular binding model.

**Fig. S3.**
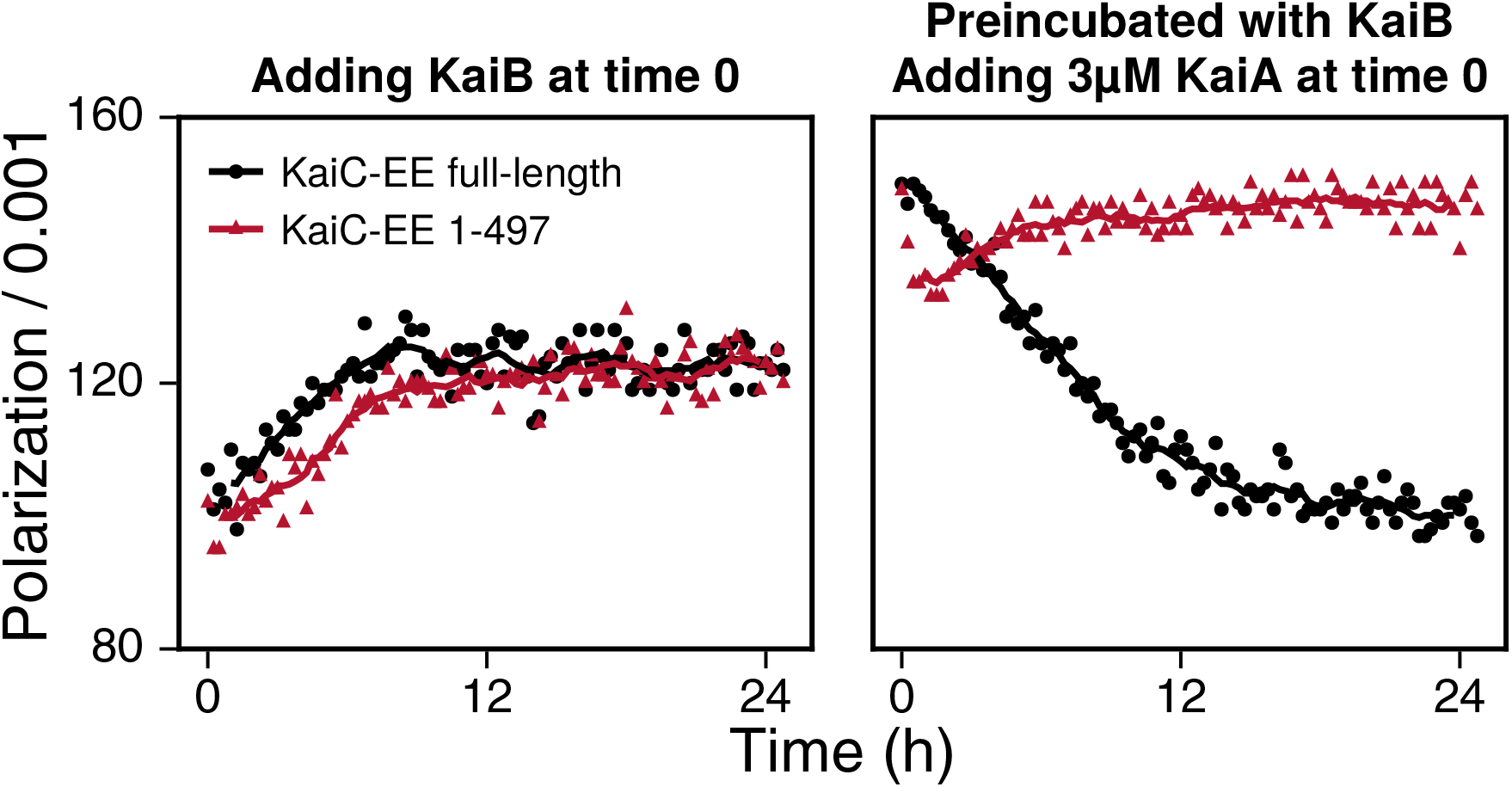
(**Left**) KaiB•KaiC complex formation kinetics measured by fluorescence polarization. Black, full-length KaiC-EE. Red, a C-terminal truncation mutant of KaiC-EE. Circles, experimental measurements. Solid lines, running average. (**Right**) Kinetic measurements of pre-formed KaiB•KaiC complex after adding 3 µM KaiA using fluorescence polarization.

**Fig. S4.**
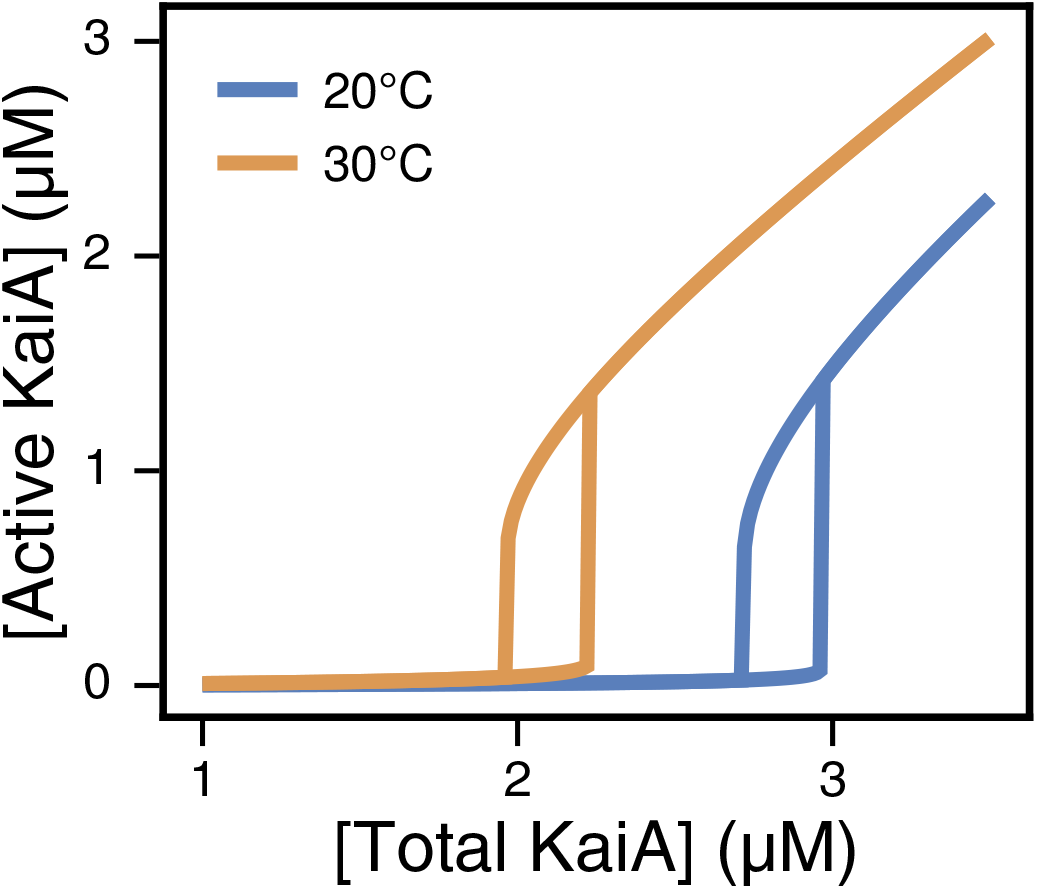
A bifurcation diagram showing the steady-state amount of active KaiA, KaiA not trapped by the KaiB•KaiC complex, predicted using the bimolecular binding model. Experimentally, the upper and lower branches can be elicited by preincubating KaiC-EE with either KaiA or KaiB. Model parameters are constrained by experimentally measured KaiB polarization kinetics.

**Fig. S5.**
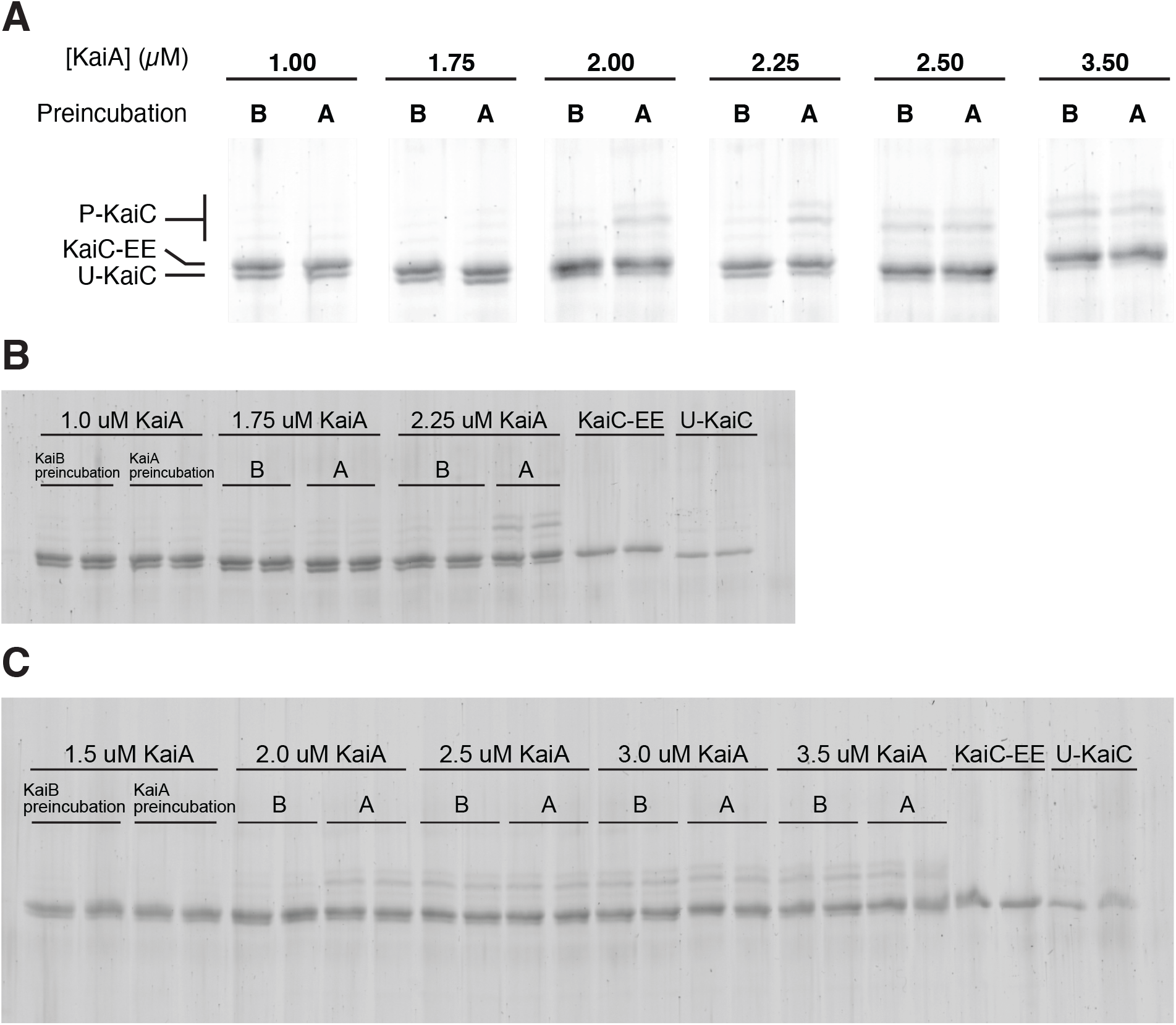
KaiA activity assay. In this assay, KaiC-EE is mixed with KaiB and different concentrations of KaiA, either with KaiB added first or KaiA added first. The amount of active KaiA in the reaction mixture is then assessed by spiking in unphosphorylated KaiC (U-KaiC) as a probe. (**A**) selected gel lanes. (**B-C**) unprocessed gel images. Wildtype KaiC is N-terminally fused with a FLAG tag to induce a gel shift compared to KaiC-EE.

**Fig. S6.**
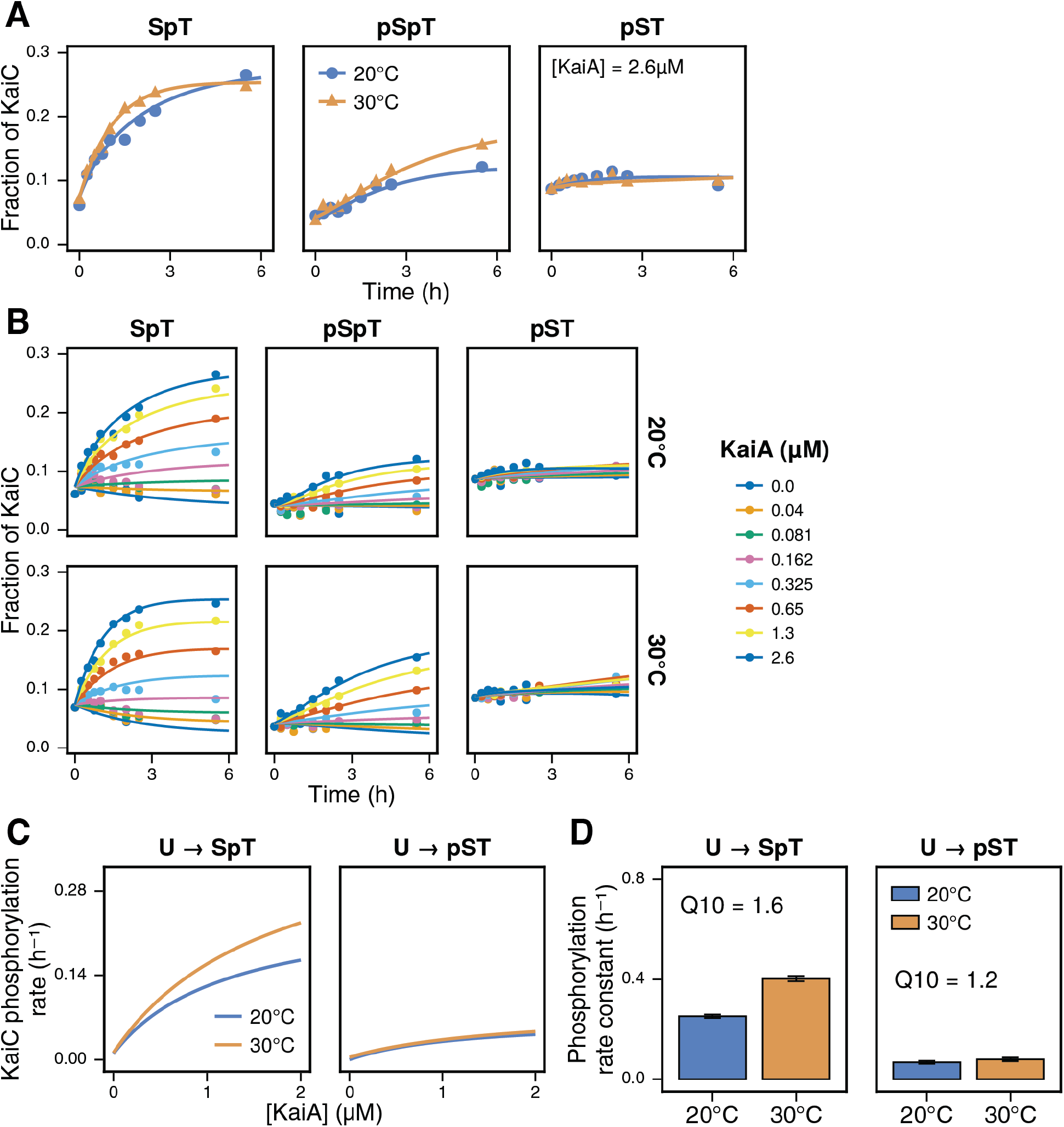
Site-specific KaiC phosphorylation kinetics starting from a dephosphorylated state. (**A-B**) Circles represent experimental measurements by SDS-PAGE; Solid lines represent fits using the model in **fig. S7**. (**A**) Representative phosphorylation kinetics in the presence of 2.6 µM KaiA (**B**) Phosphorylation kinetics under varying concentrations of KaiA. (**C-D**) Rates of selected phosphorylation steps estimated with high confidence. (**C**) Phosphorylation rates as a function of KaiA concentration. (**D**) Best-fit rate constants. Error bars, standard errors of the parameters.

**Fig. S7.**
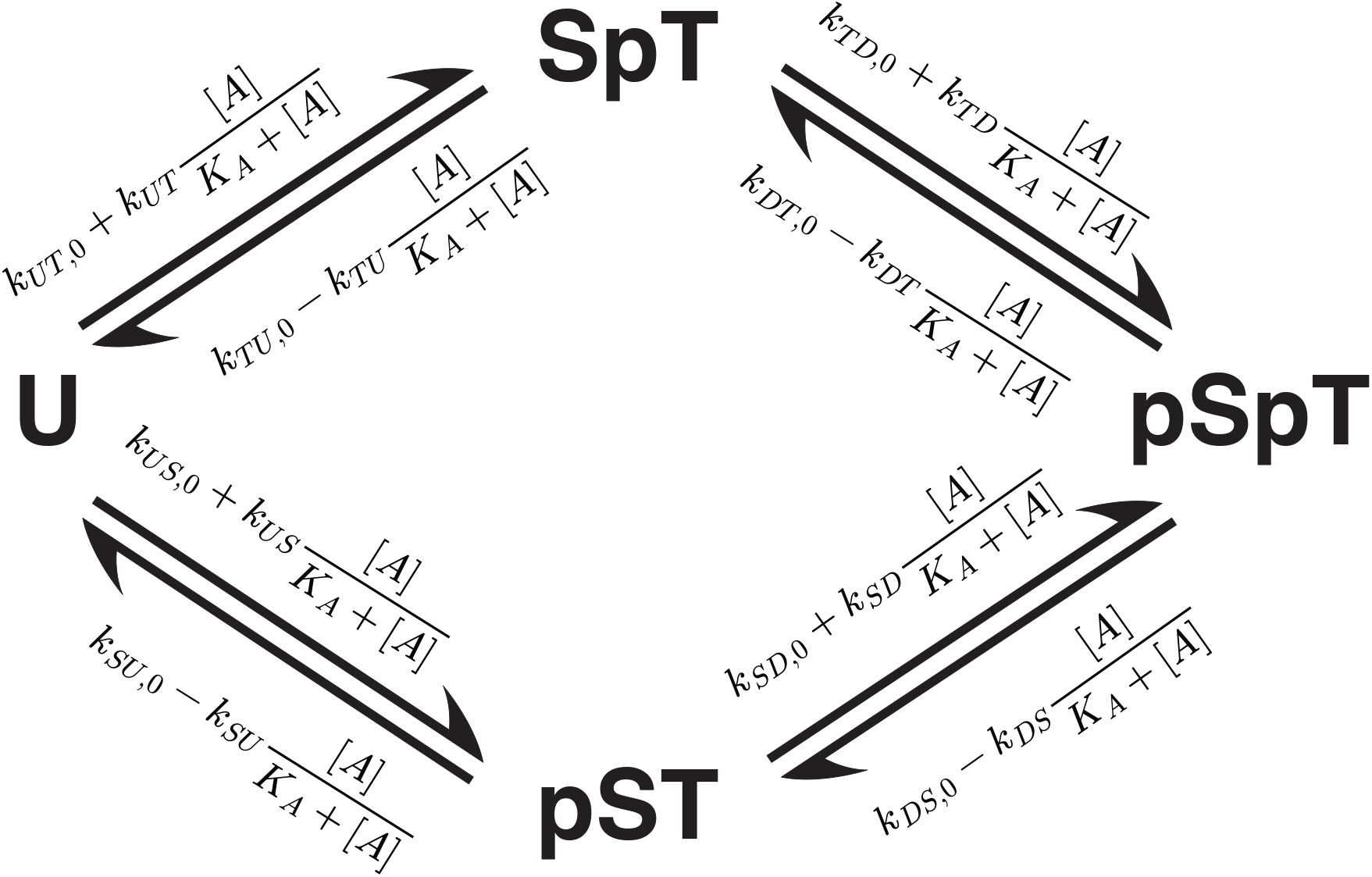
A reaction scheme of the autophosphorylation and autodephosphorylation reactions occurring at the two residues of KaiC, Ser431 and Thr432. Phosphorylation reactions are activated by KaiA, whereas dephosphorylation reactions are inhibited by KaiA. We fit this reaction model to KaiC phosphorylation and dephosphorylation data to estimate the various reaction constants. U, unphosphorylated KaiC. SpT, KaiC phosphorylated at Thr432 residue only. pSpT, KaiC doubly phosphorylated at Ser431 and Thr432. pST, KaiC phosphorylated at Ser431 only.

**Fig. S8.**
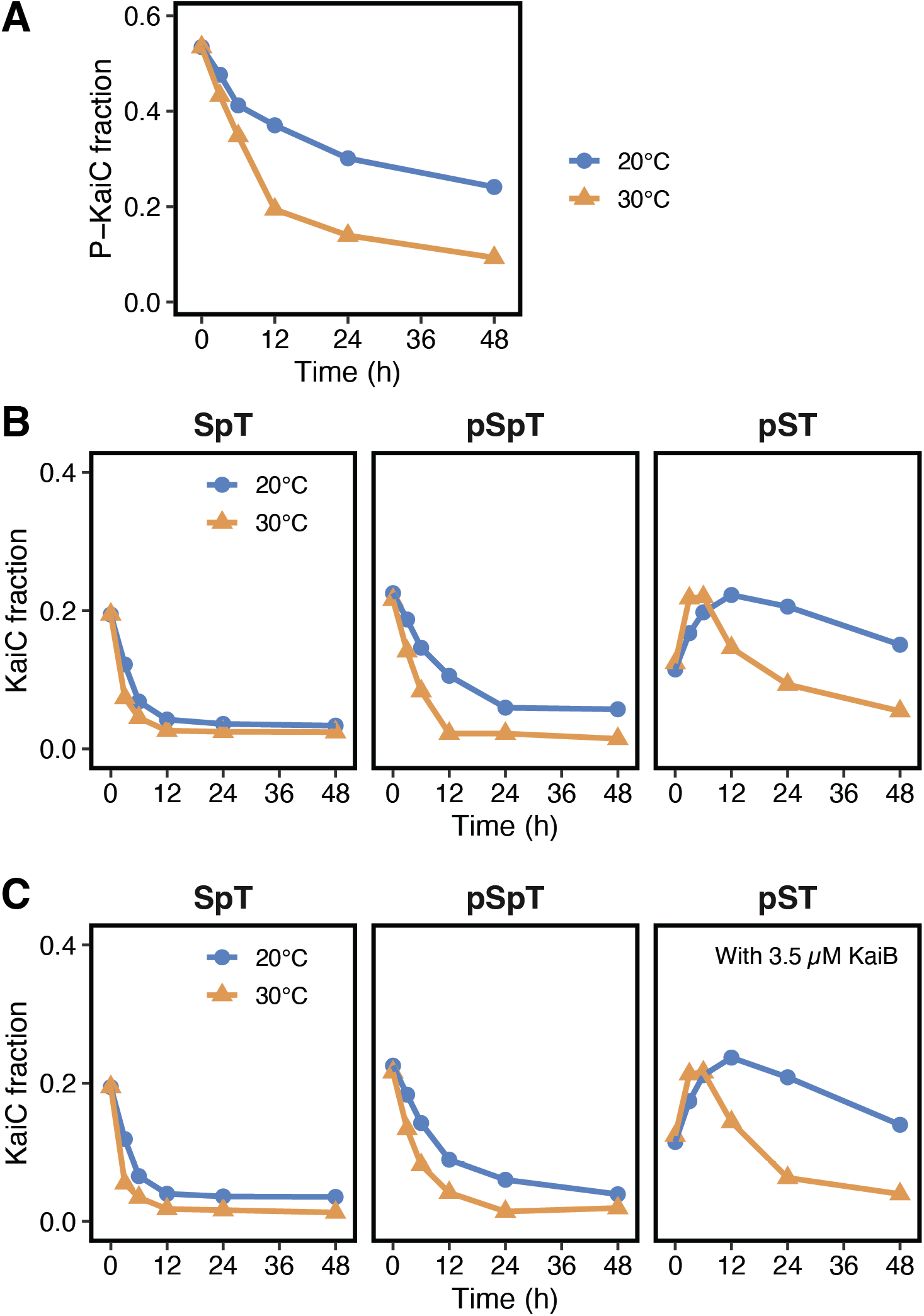
Dephosphorylation kinetics of wildtype KaiC. The reaction starts with a fresh freezer stock of purified KaiC. (**A**) Time traces of total phosphorylated KaiC (P-KaiC) in the absence of KaiB. (**B**) Site-specific kinetics in the absence of KaiB. (**C**) Site-specific kinetics the presence of KaiB.

**Fig. S9.**
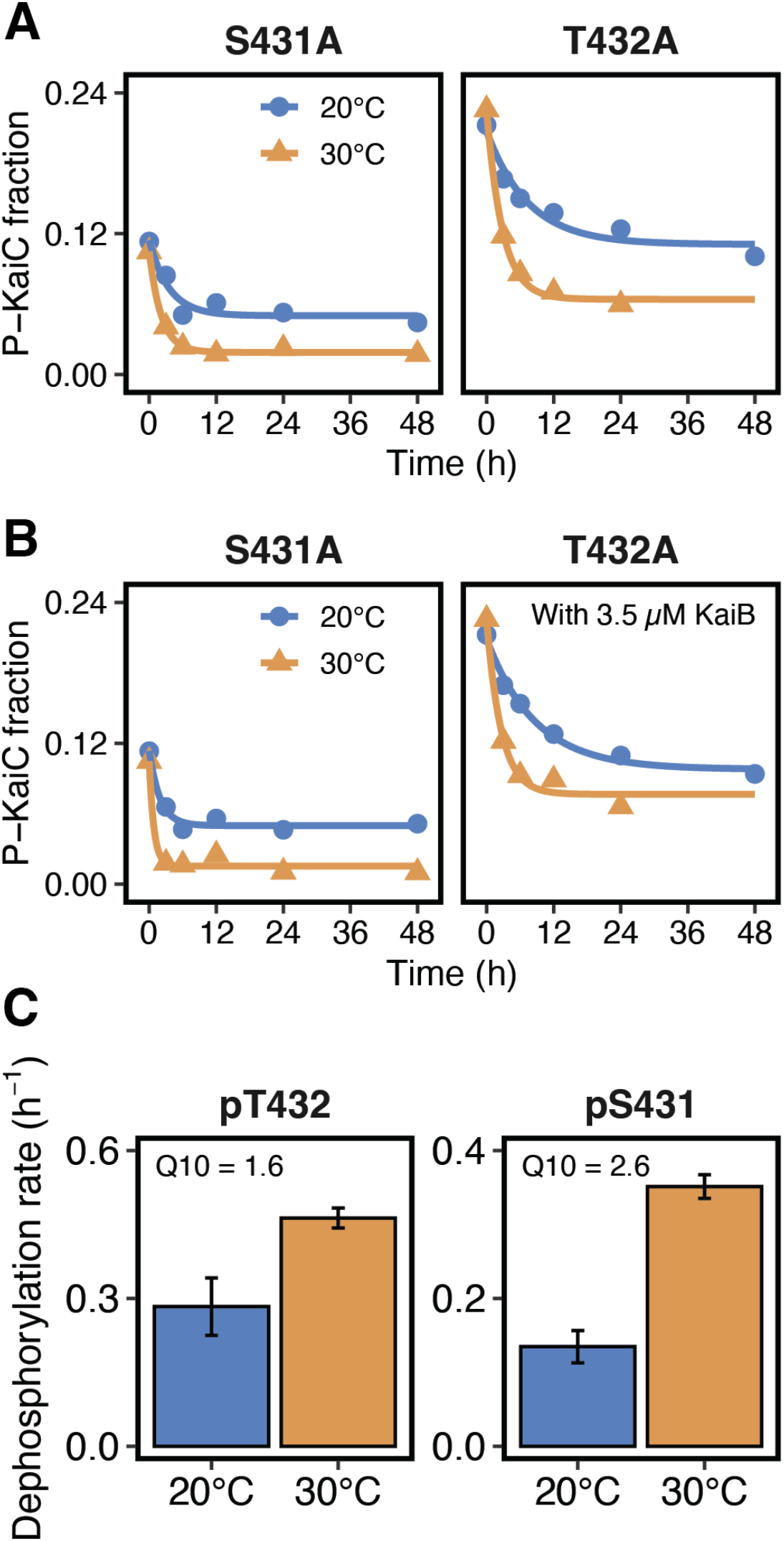
Dephosphorylation kinetics of single-site KaiC mutants S431A and T432A. Circles represent experimental measurements and solid lines represent fits to an exponential function. (**A**) In the absence of KaiB. (**B**) In the presence of KaiB. (**C**) Best-fit dephosphorylation rate constants of single-site KaiC mutants S431A (left) and T432A (right) in the absence of KaiB. Error bars, standard errors of the parameters.

**Fig. S10.**
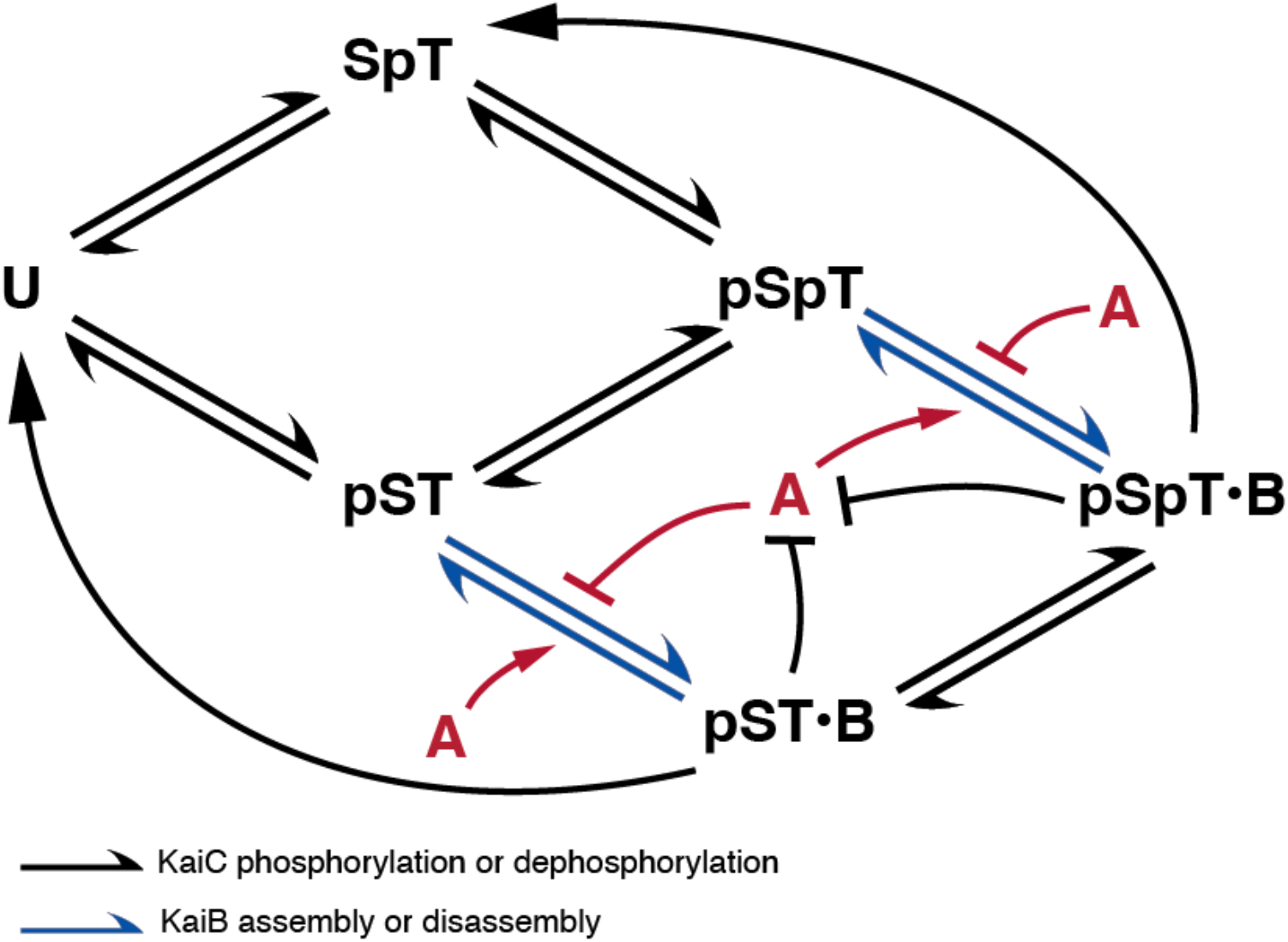
Schematic of a mathematical model of the KaiABC oscillator. This ordinary differential equation model extends the model in (*14*) by incorporating KaiA-dependent KaiB association and dissociation rates. The model is described in detail in **Supplementary Text**. Here U, SpT, pSpT, and pST represent different phosphorylation states of KaiC. U, unphosphorylated KaiC. SpT, KaiC phosphorylated at Thr432 residue only. pSpT, KaiC phosphorylated at both Ser431 and Thr432. pST, KaiC phosphorylated at Ser431 only. pSpT•B and pST•B represent pSpT-KaiC•KaiB complex and pST-KaiC•KaiB complex, respectively. A, KaiA.

**Fig. S11.**
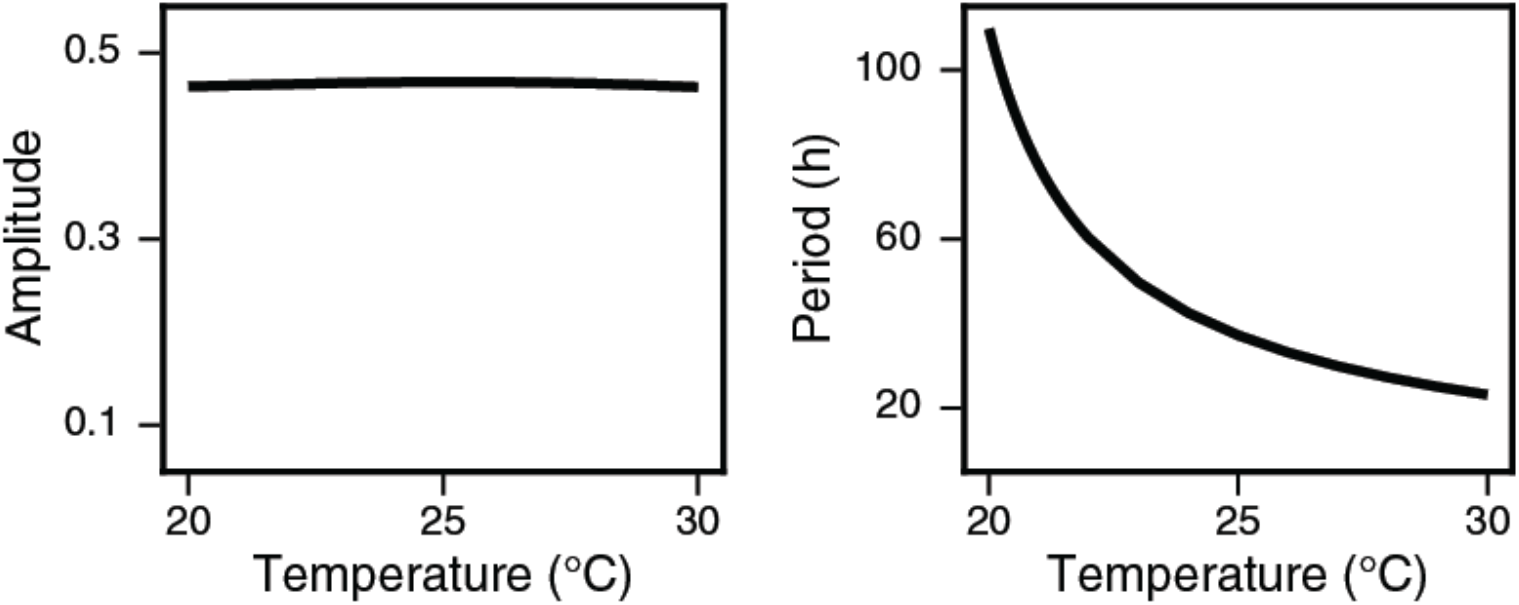
Temperature dependence of the amplitude (**Left**) and period (**Right**) of KaiABC oscillation simulated using a previously described model (*14*) which lacks the underlying bistability. Amplitude here is defined as the peak-to-trough distance of the phosphorylated KaiC fraction. Phosphorylation and dephosphorylation rate constants were scaled by preserving the original parameter set, which describes the system at 30°C, and at lower temperatures applying the Q_10_ coefficients inferred from experimental data (marked in red in **table S1**. See **Supplementary Text** for the introduction of temperature scaling in detail).

**Fig. S12.**
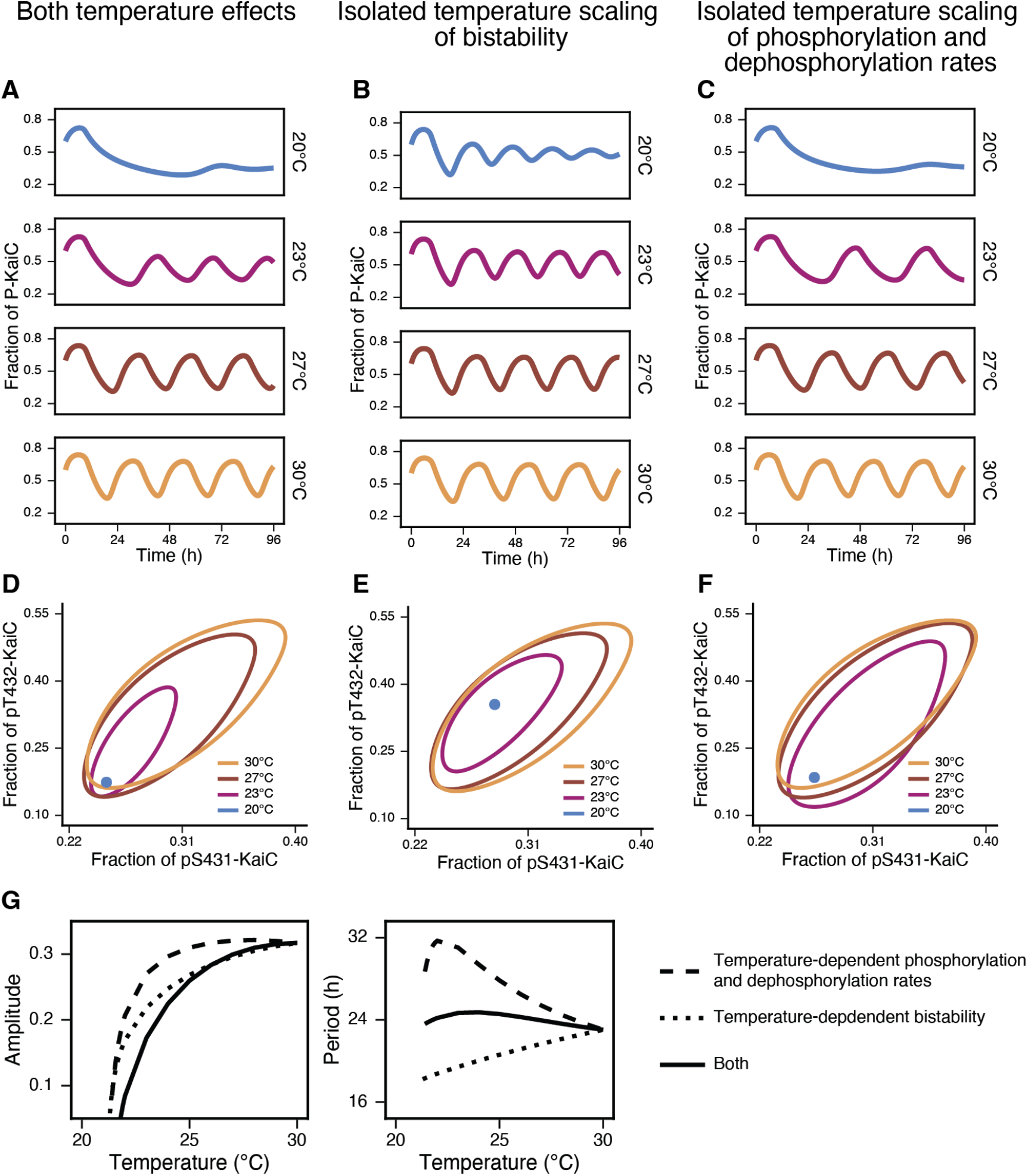
Contribution of temperature-dependent bistability scaling and reaction rate scaling to KaiABC oscillator dynamics. Three mathematical models were compared: (**A** and **D**) a full model incorporating both scaling of bistability and scaling of phosphorylation and dephosphorylation rates, further characterized in **Fig. 4F-G**, (**B** and **E**) a model incorporating scaling of bistability only, and (**C** and **F**) a model incorporating scaling of phosphorylation and dephsphorylation rates only. (**A-C**) Time traces of fractional KaiC phosphorylation. (**D-F**) Limit cycles plotted in the phosphorylation phase space of two KaiC residues, Ser431 and Thr432. (**G**) Amplitude and period changes as a function of temperature. Amplitude here is defined as the peak-to-trough distance of the phosphorylated KaiC fraction.

**Table 1.** Best-fit parameters of the KaiC phosphorylation and dephosphorylation reaction scheme in fig. S7 to experimental data. Q_10_ is defined as the ratio between the 30 °C value and the 20 °C value. Q_10_ values highlighted in red were used in the oscillator model for describing temperature scaling.

| Parameter | Unit | Description | 20°C |  | 30°C |  | Q<sub>10</sub> |
| --- | --- | --- | --- | --- | --- | --- | --- |
|  |  |  | Value | Standard Error | Value | Standard Error |  |
| $k_{UT0}$ | h<sup>-1</sup> | Baseline<br>U → SpT | 0.0101 | 0.00105 | 0.0114 | 0.00114 | 1.13 |
| $k_{UT}$ | h<sup>-1</sup> | KaiA-dependent<br>U → SpT | 0.252 | 0.0137 | 0.403 | 0.0209 | 1.60 |
| $k_{TD0}$ | h<sup>-1</sup> | Baseline<br>SpT → pSpT | 0.0340 | 0.0143 | 0.118 | 0.0191 | 3.47 |
| $k_{TD}$ | h<sup>-1</sup> | KaiA-dependent<br>SpT → pSpT | 0.748 | 0.157 | 0.0447 | 0.0773 | 0.06 |
| $k_{SD0}$ | h<sup>-1</sup> | Baseline<br>pST → pSpT | 0.0254 | 0.00286 | 0.0149 | 0.00649 | 0.59 |
| $k_{SD}$ | h<sup>-1</sup> | KaiA-dependent<br>pST → pSpT | 0.334 | 0.108 | 0.438 | 0.119 | 1.31 |
| $k_{US0}$ | h<sup>-1</sup> | Baseline<br>U → pST | ≈ 0 | 0.00176 | 0.00388 | 0.000769 | Not meaningful |
| $k_{US}$ | h<sup>-1</sup> | KaiA-dependent<br>U → pST | 0.0679 | 0.0117 | 0.0796 | 0.0146 | 1.17 |
| $k_{TU0}$ | h<sup>-1</sup> | Baseline<br>SpT → U | 0.198 | 0.0125 | 0.271 | 0.0241 | 1.37 |
| $k_{TU}$ | h<sup>-1</sup> | KaiA-dependent<br>SpT → U | 0.314 | 0.0635 | 0.365 | 0.0806 | 1.16 |
| $k_{DT0}$ | h<sup>-1</sup> | Baseline<br>pSpT → SpT | 0.0100 | 0.0105 | ≈ 0 | 0.0148 | Not meaningful |
| $k_{DT}$ | h<sup>-1</sup> | KaiA-dependent<br>pSpT → SpT | 2.00 | 0.387 | 0.401 | 0.118 | 0.20 |
| $k_{DS0}$ | h<sup>-1</sup> | Baseline<br>pSpT → pST | 0.107 | 0.00519 | 0.278 | 0.0124 | 2.60 |
| $k_{DS}$ | h<sup>-1</sup> | KaiA-dependent<br>pSpT → pST | 0.107 | 0.00520 | 0.278 | 0.0124 | 2.60 |
| $k_{SU0}$ | h<sup>-1</sup> | Baseline<br>pST → U | 0.0205 | 0.00824 | 0.119 | 0.00709 | 5.80 |
| $k_{SU}$ | h<sup>-1</sup> | KaiA-dependent<br>pST → U | ≈ 0 | 0.141 | ≈ 0 | 0.147 | Not meaningful |
| $K_A$ | μM | Michaelis-Menten<br>constant of KaiA<br>regulation | 1.25 | 0.0903 | 1.73 | 0.133 | 1.38 |

**Table 2.**
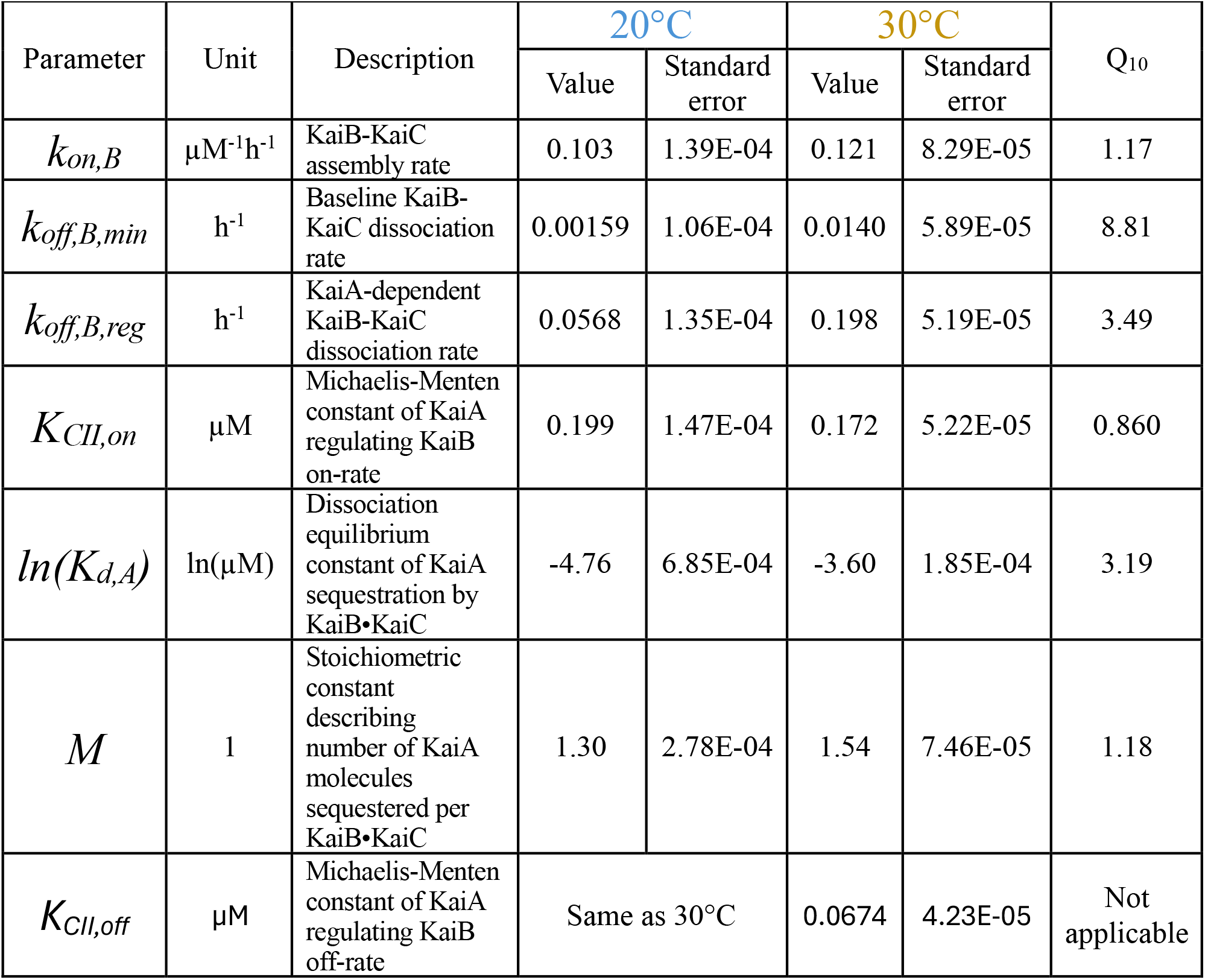
Best-fit parameters of the bimolecular binding model (**Fig. 2C**) by simultaneously fitting to the KaiB•KaiC complex formation data and KaiC phosphorylation oscillation data. The value of *K*_*CII,off*_ is fixed between the two temperatures because the KaiA regulatory effect at 20°C is too weak for this parameter, as a Michaelis-Menten constant, to be meaningfully constrained.

## References and Notes

1. C. S. Pittendrigh, in Biological rhythms. (Springer, 1981), pp. 57–80.

2. M. Elias, G. Wieczorek, S. Rosenne, D. S. Tawfik, The universality of enzymatic rate-temperature dependency. Trends Biochem Sci 39, 1–7 (2014).

3. P. L. Lakin-Thomas, S. Brody, G. G. Cote, Amplitude model for the effects of mutations and temperature on period and phase resetting of the Neurospora circadian oscillator. J Biol Rhythms 6, 281–297 (1991).

4. T. Yoshida, Y. Murayama, H. Ito, H. Kageyama, T. Kondo, Nonparametric entrainment of the in vitro circadian phosphorylation rhythm of cyanobacterial KaiC by temperature cycle. Proc Natl Acad Sci U S A 106, 1648–1653 (2009).

5. P. François, N. Despierre, E. D. Siggia, Adaptive Temperature Compensation in Circadian Oscillations. PLOS Computational Biology 8, e1002585 (2012).

6. M. Nakajima et al., Reconstitution of circadian oscillation of cyanobacterial KaiC phosphorylation in vitro. science 308, 414–415 (2005).

7. Y. Murayama et al., Low temperature nullifies the circadian clock in cyanobacteria through Hopf bifurcation. Proc Natl Acad Sci U S A 114, 5641–5646 (2017).

8. R. Pattanayek et al., Visualizing a circadian clock protein: crystal structure of KaiC and functional insights. Mol Cell 15, 375–388 (2004).

9. T. Nishiwaki-Ohkawa, Y. Kitayama, E. Ochiai, T. Kondo, Exchange of ADP with ATP in the CII ATPase domain promotes autophosphorylation of cyanobacterial clock protein KaiC. Proc Natl Acad Sci U S A 111, 4455–4460 (2014).

10. L. Hong, B. P. Vani, E. H. Thiede, M. J. Rust, A. R. Dinner, Molecular dynamics simulations of nucleotide release from the circadian clock protein KaiC reveal atomic-resolution functional insights. Proc Natl Acad Sci U S A 115, E11475–E11484 (2018).

11. M. J. Rust, J. S. Markson, W. S. Lane, D. S. Fisher, E. K. O’Shea, Ordered phosphorylation governs oscillation of a three-protein circadian clock. Science 318, 809–812 (2007).

12. T. Nishiwaki et al., A sequential program of dual phosphorylation of KaiC as a basis for circadian rhythm in cyanobacteria. EMBO J 26, 4029–4037 (2007).

13. T. Nishiwaki, T. Kondo, Circadian autodephosphorylation of cyanobacterial clock protein KaiC occurs via formation of ATP as intermediate. J Biol Chem 287, 18030–18035 (2012).

14. C. Phong, J. S. Markson, C. M. Wilhoite, M. J. Rust, Robust and tunable circadian rhythms from differentially sensitive catalytic domains. Proc Natl Acad Sci U S A 110, 1124–1129 (2013).

15. Y. G. Chang et al., Circadian rhythms. A protein fold switch joins the circadian oscillator to clock output in cyanobacteria. Science 349, 324–328 (2015).

16. Y. Furuike et al., Elucidation of master allostery essential for circadian clock oscillation in cyanobacteria. Sci Adv 8, eabm8990 (2022).

17. R. Tseng et al., Structural basis of the day-night transition in a bacterial circadian clock. Science 355, 1174–1180 (2017).

18. J. Lin, J. Chew, U. Chockanathan, M. J. Rust, Mixtures of opposing phosphorylations within hexamers precisely time feedback in the cyanobacterial circadian clock. Proc Natl Acad Sci U S A 111, E3937–3945 (2014).

19. D. Angeli, J. E. Ferrell, Jr., E. D. Sontag, Detection of multistability, bifurcations, and hysteresis in a large class of biological positive-feedback systems. Proc Natl Acad Sci U S A 101, 1822–1827 (2004).

20. J. E. Ferrell, Jr., Feedback regulation of opposing enzymes generates robust, all-or-none bistable responses. Curr Biol 18, R244–245 (2008).

21. D. Simon, A. Mukaiyama, Y. Furuike, S. Akiyama, Slow and temperature-compensated autonomous disassembly of KaiB-KaiC complex. Biophys Physicobiol 19, 1–11 (2022).

22. J. Paijmans, D. K. Lubensky, P. R. t. Wolde, A thermodynamically consistent model of the post-translational Kai circadian clock. PLOS Computational Biology 13, e1005415 (2017).

23. P. B. Kidd, M. W. Young, E. D. Siggia, Temperature compensation and temperature sensation in the circadian clock. Proceedings of the National Academy of Sciences 112, E6284–E6292 (2015).

24. H. Fu, C. Fei, Q. Ouyang, Y. Tu, Temperature compensation through kinetic regulation in biochemical oscillators. Proc Natl Acad Sci U S A 121, e2401567121 (2024).

25. I. Vakonakis, A. C. LiWang, Structure of the C-terminal domain of the clock protein KaiA in complex with a KaiC-derived peptide: implications for KaiC regulation. Proc Natl Acad Sci U S A 101, 10925–10930 (2004).

26. K. Terauchi et al., ATPase activity of KaiC determines the basic timing for circadian clock of cyanobacteria. Proc Natl Acad Sci U S A 104, 16377–16381 (2007).

27. R. Murakami et al., ATPase activity and its temperature compensation of the cyanobacterial clock protein KaiC. Genes Cells 13, 387–395 (2008).

28. Y. Murayama et al., Tracking and visualizing the circadian ticking of the cyanobacterial clock protein KaiC in solution. The EMBO journal 30, 68–78 (2011).

29. Y. Furuike, Dongyan Ouyang, Taiki Tominaga, Tatsuhito Matsuo, Atsushi Mukaiyama, Yukinobu Kawakita, Satoru Fujiwara, and Shuji Akiyama, Cross-scale analysis of temperature compensation in the cyanobacterial circadian clock system. Communications Physics 5, (2022).

30. J. E. Ferrell, Jr. et al., Simple, realistic models of complex biological processes: positive feedback and bistability in a cell fate switch and a cell cycle oscillator. FEBS Lett 583, 3999–4005 (2009).

31. R. FitzHugh, Impulses and physiological states in theoretical models of nerve membrane. Biophysical Journal 1, 445–466 (1961).

32. M. Zhou, J. K. Kim, G. W. L. Eng, D. B. Forger, D. M. Virshup, A Period2 Phosphoswitch Regulates and Temperature Compensates Circadian Period. Molecular Cell 60, 77–88 (2015).

33. A. G. Chavan et al., Reconstitution of an intact clock reveals mechanisms of circadian timekeeping. Science 374, eabd4453 (2021).

34. Y. Liu et al., Zenodo, Ed. (2026).

